# Structure basis for distinct protective mechanisms of IGHV3-23 antibodies targeting influenza hemagglutinin stem

**DOI:** 10.1101/2025.11.06.687086

**Authors:** Huibin Lv, Yang Wei Huan, Tossapol Pholcharee, Qi Wen Teo, Wenkan Liu, Akshita B. Gopal, Danbi Choi, Madison R. Ardagh, Timothy J.C. Tan, Yuanxin Sun, Arjun Mehta, Jinghang Li, Mateusz Szlembarski, Jessica J. Huang, Emily X. Ma, Lucas E. Wittenborn, Poorva Kasture, Chris K. P. Mok, Nicholas C. Wu

## Abstract

Characterization of antibodies targeting the conserved stem domain of influenza hemagglutinin (HA) is critical for developing broadly protective countermeasures against influenza virus. From a phage display human antibody library, this study discovers three group 1 HA-specific stem antibodies, namely HB31, HB34, and HB315, all of which are encoded by IGHV3-23. While HB31 and HB34 have minimal neutralization activity *in vitro*, their Fc-mediated effector functions lead to better *in vivo* protection than the potently neutralizing HB315. Consistently, cryo-EM analysis suggests that HB31 and HB34 have a higher Fc accessibility than HB315, based on their epitopes and approaching angles. HB31 and HB34 engage a pocket in the upper HA stem that is rarely targeted by known HA stem antibodies, whereas the epitope of HB315 involves the lower stem. Overall, our findings provide insights not only into the structure-function relationship of HA stem antibodies, but also into the design of next-generation influenza therapeutics.

## INTRODUCTION

Influenza remains a persistent global health threat, causing an estimated 250,000 to 500,000 deaths annually^1^. Currently, H1N1 and H3N2 subtypes of influenza A viruses as well as Victoria lineage of influenza B viruses are circulating in human^2,3^. Influenza A viruses have caused at least four major pandemics in the past, including the 1918 H1N1 “Spanish flu”, the 1957 H2N2 “Asian flu”, the 1968 H3N2 “Hong Kong flu”, and the 2009 H1N1 “swine flu”^4^. In addition, highly pathogenic avian influenza H5 and H7 subtypes are posing serious pandemic threats, with case fatality rates up to 60%^5^. In 2024, an outbreak of avian influenza H5N1 virus clade 2.3.4.4b occurred in Texas dairy cattle^6^. Since then, more than 1,000 dairy herds in the US have been infected by avian influenza H5N1 virus^7^, along with at least 70 confirmed human cases, including one fatality^8^.

Influenza A viruses encode two major surface glycoproteins, hemagglutinin (HA) and neuraminidase (NA). HA is the most abundant and immunodominant surface antigen, serving as the primary target of neutralizing antibodies elicited by infection or vaccination^9^. To date, 19 HA subtypes (H1-H19) have been identified, which are phylogenetically categorized into group 1 (H1, H2, H5, H6, H8, H9, H11, H12, H13, H16, H17, H18, H19) and group 2 (H3, H4, H7, H10, H14, H15)^10^. HA is expressed as a single-chain precursor (HA0) and assembles into a homotrimer. HA0 is then cleaved by host proteases into disulfide-linked HA1 and HA2 subunits^10^. HA contains an immunodominant head domain, which is composed entirely of HA1, resides atop the membrane-proximal stem domain, which is primarily formed by HA2. The HA head domain binds to the host sialylated receptors and is highly variable, whereas the stem domain harbors the virus-host membrane fusion machinery and is more conserved across strains and subtypes^11^. As a result, antibodies targeting the immunosubdominant HA stem often exhibit broad neutralizing activity against antigenically diverse strains and subtypes^12,13^.

Over the past two decades, many HA stem antibodies with broad cross-reactivity and protection activity have been identified^14–20^. Analysis of the HA stem antibody repertoires has uncovered several recurring sequence features, mainly defined by immunoglobulin heavy variable (*IGHV*) gene usage and sequence motifs within the complementarity-determining region (CDR) H3. Notable examples include *IGHV6-1* with an *IGHD3-3*-encoded CDR H3 FG[I/V/L] motif^15,21,22^, *VH1-18* with a QxxV motif^21,23^, and *IGHD3-9*-encoded LxYFxWL motif^24,25^. Two major neutralizing epitopes were discovered for the group 1 HA stem, namely the central stem epitope and the anchor epitope^26–30^. While central stem antibodies often broadly recognize multiple subtypes^26–28^, anchor antibodies are usually more subtype-restricted^29,30^. Structural studies have revealed that central stem antibodies frequently target five hydrophobic pockets within the stem domain using aromatic and hydrophobic side chains^10,31^. The discovery and characterization of HA stem antibodies have motivated the development of broadly protective influenza vaccines^32–34^, some of which have shown promising results in phase I clinical trials^35–38^. HA stem antibodies have also guided structure-based design of proteins^39–43^, peptides^31,44^, and small molecules^45–47^ with neutralization activity.

In this study, we identified three *IGHV3-23* HA stem antibodies, HB31, HB34, and HB315, from a phage display human antibody library that was previously constructed from 245 healthy donors^48^. All three antibodies bound to HAs from multiple group 1 subtypes, including one from the H5N1 clade 2.3.4.4b. HB31 and HB34 primarily relied on Fc-mediated effector functions for their in vivo protection activity, whereas HB315 did not. Such a difference can be attributed to their epitopes and the approach angles revealed by cryo-EM analysis. Throughout this study, HA residue positions are numbered according to the H3 HA reference, and antibody residues are numbered using Kabat nomenclature.

## RESULTS

### Discovery of HA stem antibodies by phage display and long-read sequencing

We previously constructed a phagemid antibody library using the peripheral blood mononuclear cells (PBMCs) from 245 healthy individuals^48^. Here, this antibody library was displayed on M13 KO7 ΔpIII hyperphage (hyperphage) and CM13 interference-resistant helper phage (CM phage) (**Figure 1A**). These two phage display antibody libraries were subjected to three rounds of panning against an HA stem construct derived from the A/Brisbane/59/2007 (H1N1) HA (hereafter referred to as the H1 stem)^34^ (**Figure 1A**). For both phage display antibody libraries, the titer increased from round to round, indicating enrichment of phage clones encoding antibodies that bound to the H1 stem (**Figure 1B**). The occurrence frequencies of individual clones in the input phagemid antibody library and post-round three selection phage display antibody libraries was analyzed by PacBio long-read sequencing.

**Figure 1.**
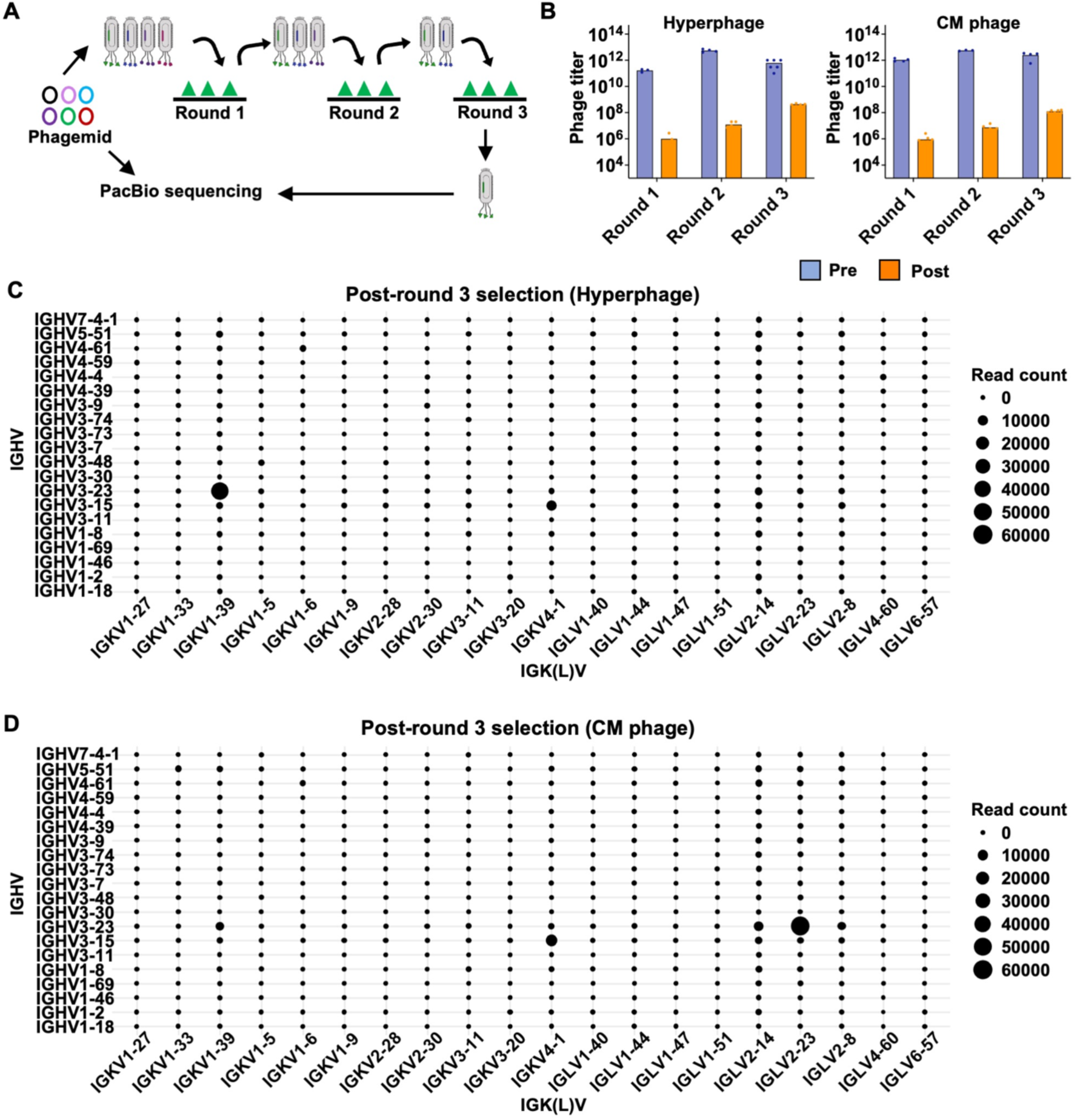
Antibody screening against HA stem by phage display. **(A)** Three rounds of panning were performed against the HA stem, followed by PacBio sequencing of both the phagemid antibody library before panning and after three rounds of panning. **(B)** Phage titers were measured before (Pre, blue) and after (Post, orange) each round of panning. **(C-D)** Distribution of *IGHV/IGK(L)V* pairings following three rounds of panning of the **(C)** Hyperphage antibody library or the **(D)** CM13 phage antibody library. Point size reflects the sequencing read count for each *IGHV/IGK(L)V* combination.

While the occurrence frequencies of different *IGHV/IGK(L)V* gene pairs were quite evenly distributed in the input phagemid antibody library **(Figure S1)**, certain *IGHV/IGK(L)V* gene pairs became enriched post-selection (**Figure 1C-1D**). After three rounds of selection, *IGHV3-23/IGLV2-23* and *IGHV3-23/IGKV1-39* had the highest occurrence frequency in the CM phage and hyperphage libraries, respectively. Our downstream analyses focused on two representative antibodies from these *V* gene families, namely HB34 (*IGHV3-23/IGLV2-23*) from the CM phage library and HB315 (*IGHV3-23/IGKV1-39*) from the hyperphage library (**Figure 2A**). We also included another antibody, HB31 (*IGHV3-23/IGLV2-8*), which had the same heavy chain variable domain sequence as HB34 but a different light chain (**Figure 2A**). HB31, HB34, and HB315 all reached high occurrence frequencies after three rounds of selection and were validated by ELISA to bind H1 stem (**Figure 2B**).

**Figure 2.**
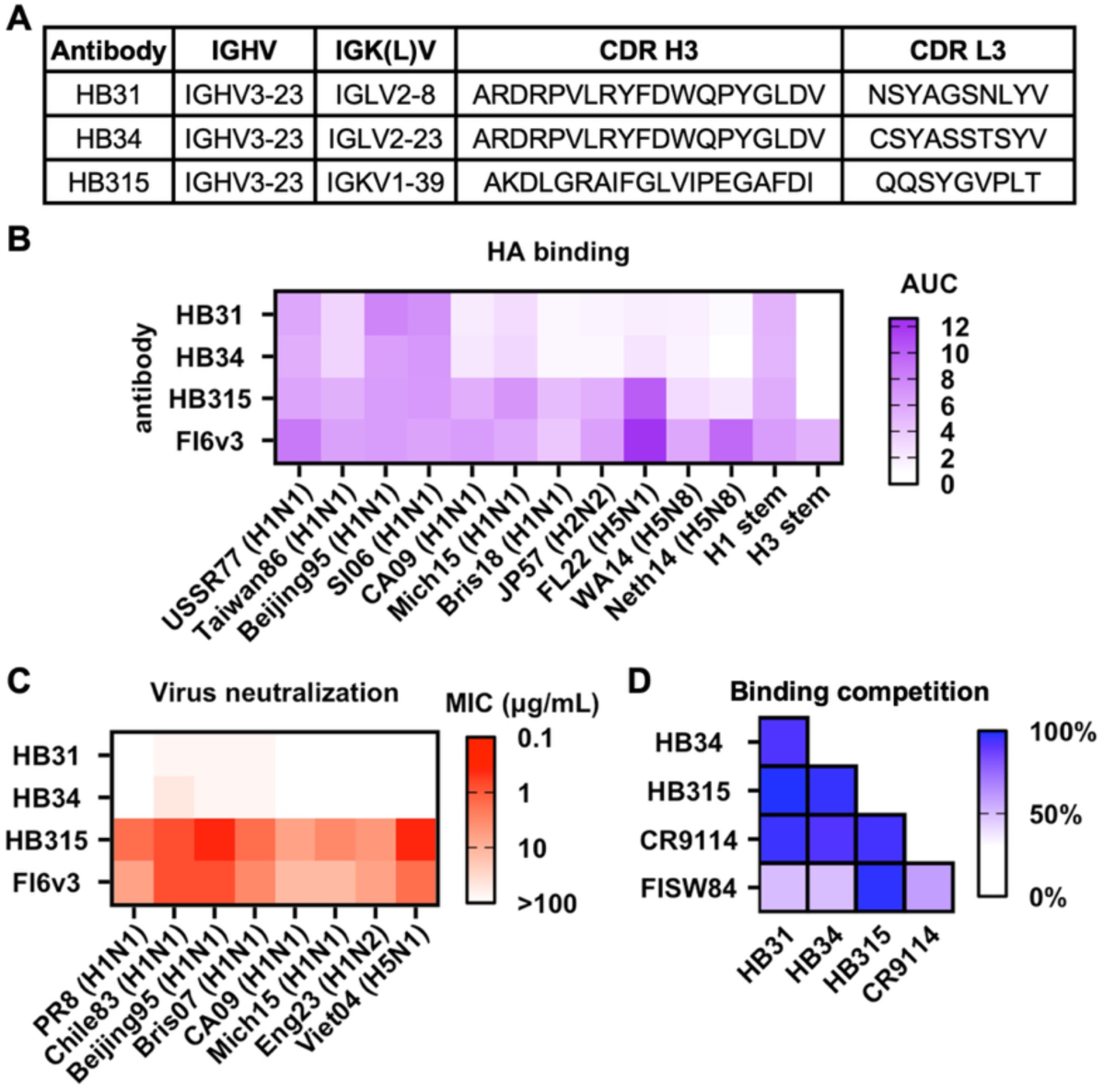
Binding and neutralization activity of HB31, HB34, and HB315. **(A)** Sequence information of three HA stem antibodies. **(B)** Binding activity of the indicated antibodies in IgG format to various HA proteins was assessed by ELISA and quantified as the area under the curve (AUC). **(C)** Neutralization activity of three HA stem antibodies was determined by microneutralization assay. FI6v3, which is a human HA stem antibody^16^, served as a positive control. **(B-C)** Strain names are abbreviated as follows: PR8: A/Puerto Rico/8/1934 (H1N1), USSR77: A/USSR/90/1977 (H1N1), Chile83: A/Chile/1/1983 (H1N1), Taiwan86: A/Taiwan/01/1986 (H1N1), Beijing95: A/Beijing/262/1995 (H1N1), SI06: A/Solomon Island/3/2006 (H1N1), Bris07: A/Brisbane/59/2007 (H1N1), CA09: A/California/07/09 (H1N1), Mich15: A/Michigan/45/2015 (H1N1), Bris18: A/Brisbane/02/2018 (H1N1), Eng23: A/England/234600203/2023 (H1N2), JP57: A/Japan/305/1957 (H2N2), Vie04: A/Vietnam/1203/2004 (H5N1), FL22: A/bald eagle/Florida/W22–134-OP/2022 (H5N1), WA14: A/northern pintail/WA/40964/2014 (H5N8), and Neth14: A/chicken/Netherlands/14015531/2014 (H5N8). **(D)** Competition among the indicated antibodies for binding to HA H1N1 A/Michigan/45/2015 (H1N1) was measured by biolayer interferometry (BLI). All antibodies were in Fab format. Percentage of binding competition is shown as a heatmap.

### HB315, HB31, and HB34 are group 1 HA-specific broadly neutralization antibodies

To evaluate the cross-reactivity breadth of HB31, HB34, and HB315, we assessed their binding activity against a panel of HAs from different subtypes. The three antibodies bound to all tested HAs from human H1N1 strains isolated between 1977 and 2018. Nevertheless, HB31 and HB34 exhibited weaker binding activity than HB315 against the HAs from post-2009 pandemic H1N1 strains, including A/California/04/2009 (H1N1), A/Michigan/45/2015 (H1N1), and A/Brisbane/02/2018 (H1N1) (**Figure 2B**). Similarly, the binding activity of HB31 and HB34 to HAs from H2 and H5 subtypes was weaker than HB315 (**Figure 2B**). None of the three antibodies bound to a stabilized stem construct derived from H3N2 A/Finland/486/2004 HA (group 2 HA)^49^, indicating that their specificity was specific to group 1 HA (**Figure 2B**).

We further tested the neutralizing activity of HB31, HB34, and HB315 against a panel of human H1N1 strains, avian A/Vietnam/1203/2004 (H5N1), and swine A/England/234600203/2023 (H1N2). Among the three antibodies, HB315 exhibited higher neutralization activity across all tested viruses (**Figure 2C**). On the contrary, HB31 and HB34 showed limited neutralization, with weak activity against A/Chile/1/83 (H1N1), A/Beijing/262/95 (H1N1), and A/Brisbane/59/2007 (H1N1), and no detectable activity against the remaining strains, even at 100 μg/mL (**Figure 2C**). Collectively, HB315 demonstrated superior neutralization potency and breadth compared to HB31 and HB34.

### HB315 has a different antibody competition profile from HB31 and HB34

To probe the epitopes of HB31, HB34, and HB315, we performed a binding competition assay. This binding competition assay also included FISW84^29^ and CR9114^14^ as controls, which are prototypic antibodies to the anchor and central stem epitopes, respectively. HB31 and HB34 exhibited a similar competition, showing 80-90% competition with HB315 and CR9114 but only ∼50% competition with the anchor antibody FISW84 (**Figure 2D and Figure S2A-S2D)**. By contrast, HB315 competed strongly with all tested antibodies, including FISW84 (**Figure 2D and Figure S2A-S2D)**. These results suggested that HB31 and HB34 likely targeted the central stem epitope, similar to CR9114, whereas the epitope of HB315 likely spanned both the anchor and central stem epitopes.

### HB31, HB34, and HB315 confer *in vivo* protection via different mechanisms

To test the *in vivo* protection activity of HB31, HB34, and HB315, all three antibodies were administrated prophylactically prior to a lethal challenge with the A/Puerto Rico/8/1934 (PR8, H1N1) virus. According to the weight loss profiles (**Figure 3A, 3C, and 3E)**, survival analysis (**Figure 3B, 3D, and 3F)**, and viral lung titer at day 3 post-infection (**Figure 3L**), all three antibodies conferred protection against PR8. Nevertheless, the protection activity of HB31 and HB34 were stronger than HB315. With 1 mg/kg of antibodies, the survival rates of mice treated with HB31, HB34, and HB315 were 80% (4/5), 80% (4/5), and 20% (1/5), respectively (**Figure 3C-D**). At 0.3 mg/kg, none of the mice treated with HB315 survived, whereas the survival rates of the mice treated with HB31 and HB34 were 60% (3/5) and 40% (2/5), respectively (**Figure 3E-F**). Furthermore, we tested the prophylactic protection activity of these antibodies at 5 mg/kg against a lethal challenge with A/Vietnam/1203/2004 (H5N1). While all the three antibodies reduced the viral lung titer on day 3 post-infection (**Figure 3K**), only HB31 conferred complete protection, resulting in 100% survival rate of H5N1-infected mice (**Figure 3H**). On the other hand, HB34 and HB315 conferred only partial protection, with survival rates of 60% (3/5) and 40% (2/5), respectively (**Figure 3G-3H**). Together, these results revealed an inverse relationship between neutralization potency and protection activity, with HB31 and HB34 exhibiting the weaker neutralization potency *in vitro* but conferring the stronger protection activity *in vivo* than HB315.

**Figure 3.**
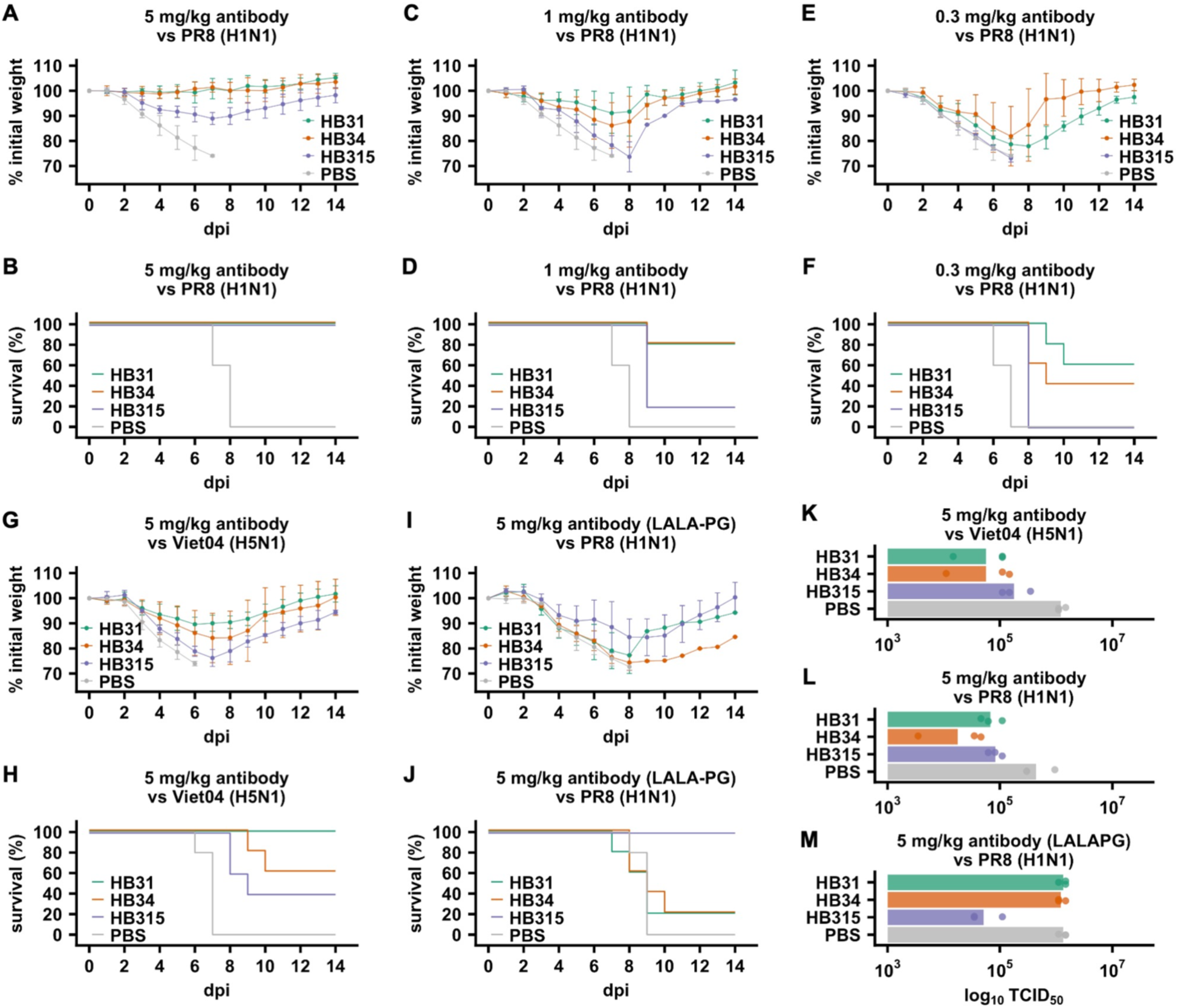
*In vivo* protection activity of HB31, HB34, and HB315. Female BALB/c mice at 6 weeks old were injected intraperitoneally with the indicated antibodies at the indicated dose 4 hours prior to challenge with **(A-F, I, J, L, M)** 5× LD_50_ of A/Puerto Rico/8/1934 (PR8, H1N1) or **(G, H, K)** 5× LD_50_ of recombinant PR8 virus carrying the HA from A/Vietnam/1203/2004 (H5N1). **(A, C, E, G, I)** The mean percentage of body weight change post-infection is shown (n = 5). The humane endpoint was defined as a weight loss of 25% from initial weight on day 0. **(B, D, F, H, J)** Kaplan-Meier survival curves are shown (n = 5). **(K, L, M)** Lung viral titers on day 3 post-infection are shown (n = 3).

Previous studies have shown that non-neutralizing HA stem antibodies can still confer protection *in vivo* due to Fc-mediated effector functions^50,51^. To test if Fc-mediated effect functions of HB31, HB34, and HB315 contributed to their protection activity *in vivo,* we generated a LALA-PG variant for each of the three antibodies to eliminate their Fc effector functions^52^ **(Figure S3)**. While all mice treated with 5 mg/kg of HB315-LALA-PG were protected against PR8, the survival rate of mice treated with 5 mg/kg of HB31-LALA-PG or HB34-LALA-PG were only at 20% (1/5) (**Figure 3I-J**). Additionally, mice treated with HB31-LALA-PG or HB34-LALA-PG had similar viral lung titer at day 3 post-infection as those without antibody treatment (**Figure 3M**). In comparison, the viral lung titer at day 3 post-infection in mice treated with HB315-LALA-PG was reduced by more than 10-fold. These results suggested that Fc effector functions were more important for the *in vivo* protection activity of HB31 and HB34 than for that of HB315. Consistently, HB31 and HB34 elicited stronger antibody-dependent cell-mediated cytotoxicity (ADCC) activity than HB315 in an *in vitro* reporter assay **(Figure S3)**.

### The epitope of HB315 overlaps with but is distinct from that of HB31 and HB34

To understand the biophysical basis of the distinct protection mechanisms among HB31, HB34, and HB315, we determined the cryo-EM structure of the fragment antigen-binding (Fab) of HB31, HB34 and HB315 in complex with A/Solomon Islands/3/2006 (H1N1) HA to a resolution of 2.76 Å, 2.64 Å, and 2.67 Å, respectively (**Figure 4A-C and Table S1**). All the three antibodies interacted with HA exclusively through their heavy chains (**Figure S4A-C**). HB31 and HB34 engaged HA stem with a horizontal approaching angle, whereas HB315 engaged HA stem with an upward approaching angle (**Figure 4A-D**). This upward approaching angle of HB315 likely limited the accessibility of its Fc to effector cells, explaining its lower ADCC activity than HB31 and HB34 (**Figure S3A-B**), as well as its minimal reliance on Fc-mediated effector functions for protection activity *in vivo* (**Figure 3I and 3M**). Similarly, most anchor antibodies, which also have an upward approaching angle, lack ADCC activity^29^.

**Figure 4.**
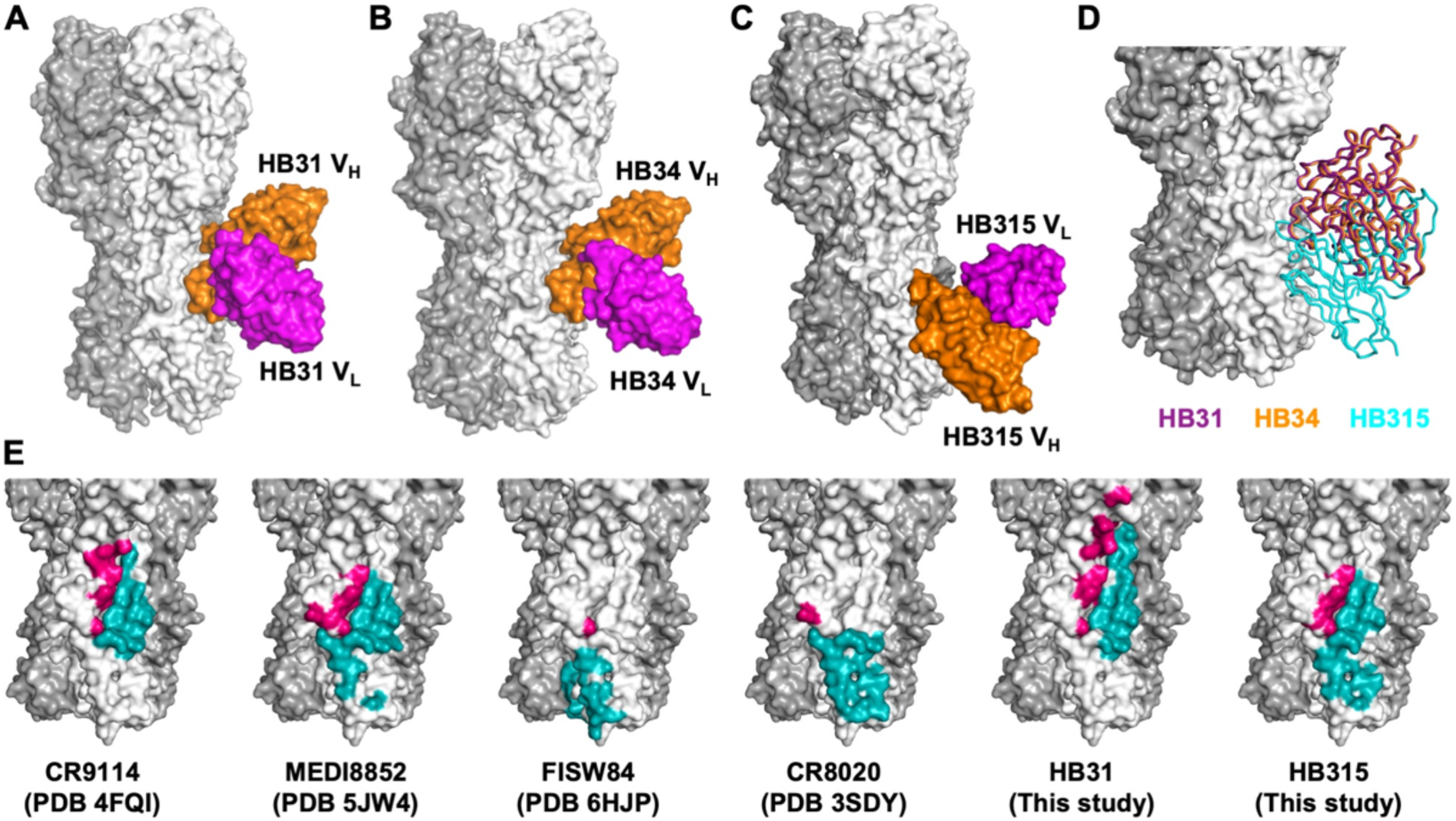
Epitopes and approaching angles of HB31, HB34, and HB315. (A-C) Cryo-EM structures of the indicated antibodies in complex with A/Solomon Island/3/06 (H1N1) HA. Antibodies and HA are shown in surface representation. HA is in white. Heavy chain variable domain (V_H_) and light chain variable domain (V_L_) are in orange and magenta, respectively. **(D)** The approaching angles of HB31 (magenta), HB34 (orange), and HB315 (blue) are compared. **(E)** Epitopes of CR9114 (PDB 4FQI)^14^, MEDI8852 (PDB 5JW4)^15^, FISW84 (PDB 6HJP)^53^, CR8020 (PDB 3SDY)^19^, HB31, and HB315 are shown on HA surface. HA1 and HA2 contacts are colored in pink and blue, respectively.

The epitope of HB315 closely resembled that of the cross-group neutralizing antibody MEDI8852^15^ (**Figure 4E**). HB315 had an *IGHD3-3*-encoded FGL motif at the tip of CDR H3 that was chemically similar to the YGV motif of MEDI8852 at equivalent positions **(Figure S4F and Figure S5B)**. However, HB315 engaged the lower HA stem more extensively compared to MEDI8852 (**Figure 4E**), mainly due to the H-bonds between its CDR H2 and the side chains of D19_HA2_, S32_HA2_, Y34_HA2_, and K153_HA2_ (**Figure S4D-E**). Notably, the epitope of HB315 had a large overlap with that of the anchor antibody FISW84^53^ and those of the group 2 HA-specific lower stem antibodies CR8020^19^ (**Figure 4E**). This observation explained the strong binding competition between HB315 and FISW84.

### HB31 and HB34 extend binding to a new pocket

HA stem contains five hydrophobic pockets that are often targeted by the central stem antibodies (**Figure 5A**)^10^. Similarly, HB31 also interacted with three of these five hydrophobic pockets (**Figure 5B**) using a *IGHD3-9*-encoded LxYFxW motif in the CDR H3 (**Figure 5C and Figure S5A)**, which is a recurring sequence feature of central stem antibodies^24,25^. HB31 further engaged another pocket in the upper stem (hereinafter referred to as pocket 6) (**Figure 5B**). This pocket was rarely targeted by the central stem antibodies. For example, the *IGHD3-9*-encoded central stem antibodies, FI6v3 and S9-3-37, did not engage pocket 6 **(Figure S6)**, despite targeting the central stem epitope using the LxYFxW motif akin to HB31 (**Figure 5C**). Nevertheless, the periphery of pocket 6 is targeted by two recently reported antibodies, 05.GC.w13.01 and 05.GC.w13.02^54^, via H-bonds (**Figure 5B and 5D**). In comparison, the interaction between HB31 and pocket 6 was more substantial. HB31 filled pocket 6 by V_H_ R55 and H-bonded with the backbone carbonyls of L292_HA1_ and S290_HA1_ deep inside pocket 6 (**Figure 5D**). The inferred germline of HB31 had a Gly at V_H_ residue 55 **(Figure S8A)**. Reverting V_H_ R55 to V_H_ G55 in HB31 weakened the binding affinity (K_D_) against A/Texas/37/2024 (H5N1) HA by more than 2-fold (**Figure 5E**), and against H1 stem by more than 30% **(Figure S8B)**, showing that engagement of pocket 6 strengthens the binding of HB31 to HA stem. Notably, these structural observations regarding HB31 also applied to HB34, which had an almost identical binding mode as HB31. By targeting the upper stem, HB31 and HB34 were likely to have high Fc accessibility, explaining their strong reliance on Fc-mediated effector functions for protection activity *in vivo*.

**Figure 5.**
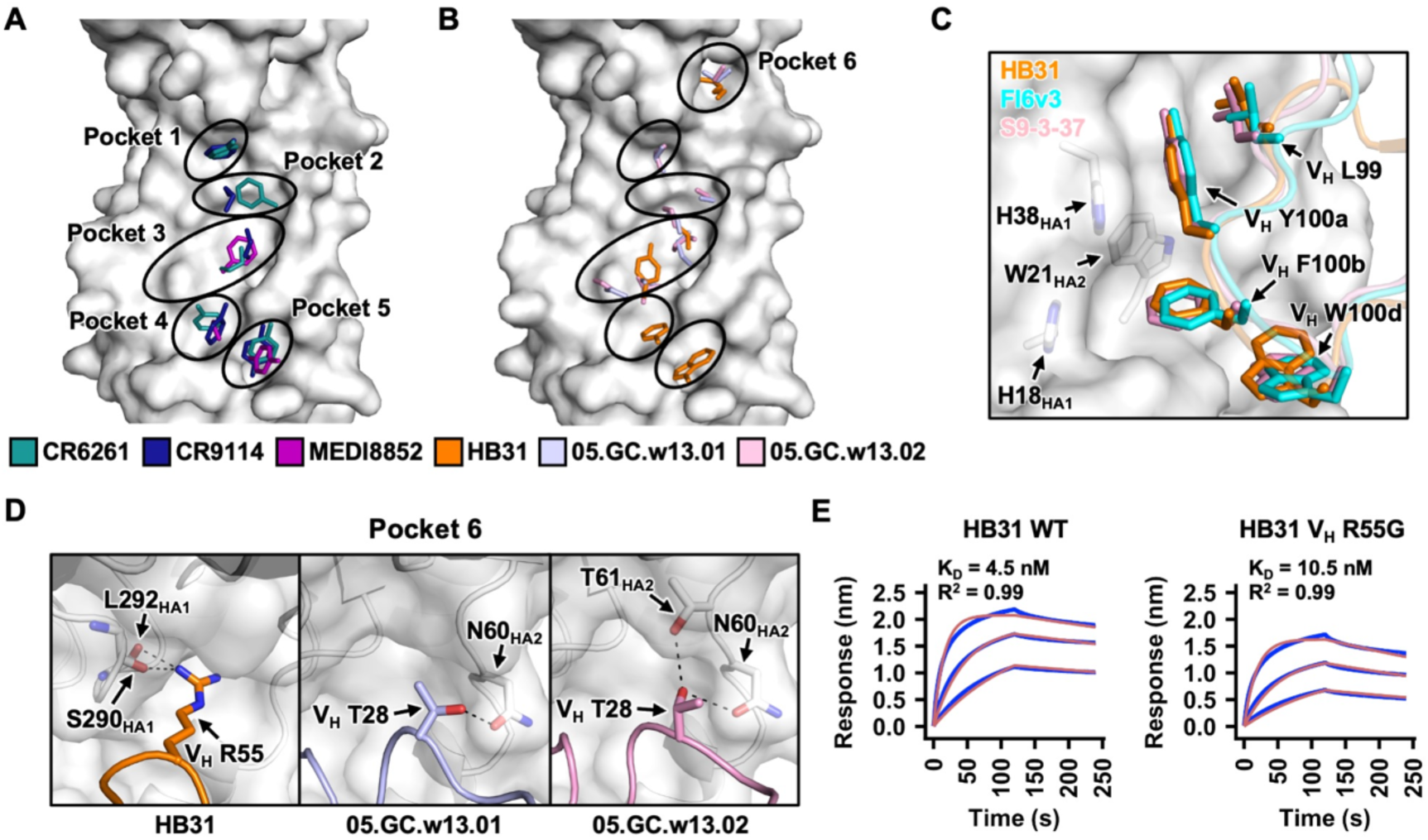
Structural features of HB31. **(A)** Five pockets in the HA stem that are targeted by known central stem antibodies are indicated on HA (white surface)^26^. Side chains of paratope residues in human HA stem antibodies CR6261 (PDB 3GBM)^26^, CR9114 (PDB 4FQI)^14^, and MEDI8852 (PDB 5JW4)^15^ that interact with these five pockets are shown as sticks. **(B)** Side chains of paratope residues in human HA stem antibodies HB31, 05.GC.w13.01 (PDB 8TXP)^54^ and 05.GC.w13.02 (PDB 8TXM)^54^ pockets 1-6 in the HA stem are shown as sticks. **(C)** Interactions between HA stem and the *IGHD3-9*-encoded LxYFxW motifs in HB31, FI6v3 (PDB 3ZTN)^16^, and S9-3-37 (PDB 6E3H)^24^ are shown. Their CDR H3s are shown as cartoon representation with the side chains of the LxYFxW motifs as sticks. HA is shown as white semitransparent surface with the side chains of key epitope residues as sticks. **(D)** Interactions of pocket 6 with HB31, 05.GC.w13.01 (PDB 8TXP)^54^ and 05.GC.w13.02 (PDB 8TXM)^54^ are compared. CDR H2s are shown as cartoon representation. Key interacting residues are shown as sticks. **(E)** Binding kinetics of HB31 wild-type (WT) and V_H_ R55G germline revertant against A/Texas/37/2024 (H5N1) HA were measured by biolayer interferometry (BLI). Y-axis represents the response. Blue and red lines represent the response curve and the 1:1 binding model, respectively. Binding kinetics were measured for 300 nM, 100 nM, and 33 nM of Fab. The dissociation constants (K_D_) and the goodness-of-fit values (R^2^) are shown.

## DISCUSSION

To date, there are three major HA stem epitopes being characterized, the central stem epitope^10^, the group 1 HA-specific anchor epitope^29,53^, and the group 2 HA-specific lower stem epitope^18,19^. In this study, we showed that the HB315 epitope has large overlap with both the central stem and the lower stem epitopes, suggesting that the three major HA stem epitopes may not be as discrete as previously thought. In addition, HB31 and HB34 target a pocket in the upper stem (pocket 6) that is rarely engaged by other known HA stem antibodies, further underscoring the epitope diversity of HA stem antibodies. While peptides and small molecules targeting pockets 1-5 of HA stem can achieve neutralization activity in the nM range^31,46^, extending those designs to target pocket 6 may further improve their potency. Together, our findings not only refine the knowledge of HA stem antigenicity, but also provide structural insights into the development of next-generation therapeutics.

Although HB31, HB34, and HB315 are all encoded by *IGHV3-23*, they contain different *IGHD*-encoded motifs that are common in the CDR H3 of HA stem antibodies. HB31 and HB34 contain an *IGHD3-9*-encoded LxYFxW motif, whereas HB315 contains an *IGHD3-3*-encoded FGL motif. Consequently, the binding modes of HB31 and HB34 are different from that of HB315. This observation demonstrates that *IGHV3-23* provides a versatile framework for HA stem antibodies to adopt different *IGHD*-driven binding modes. It also suggests that at least some *IGHD* genes have a larger influence than *IGHV* genes on the binding mode of HA stem antibodies. Consistently, while *IGHV1-69* HA stem antibodies with diverse CDR H3 sequences often have similar binding modes^20^, the presence of an *IGHD3-9*-encoded LxYFxW motif changes their binding mode to become similar to that of other HA stem antibodies that possess the same motif^55^. As *IGHD*-encoded motifs have received increasing attention as recurring sequence features against different pathogens^55–58^, future studies should explore whether HA stem antibodies with *IGHD3-9*-encoded LxYFxW motif and *IGHD3-3*-encoded FG[I/V/L] motif always result in converging binding modes regardless of *IGHV* gene usage.

A notable observation in this study is that HB31 and HB34, which have minimal neutralization activity *in vitro*, can confer similar, if not superior, protection activity *in vivo* than the potently neutralizing HB315. While neutralizing antibodies are traditionally viewed as the primary mediators of antiviral protection, growing evidence indicates that non-neutralizing antibodies can also confer strong *in vivo* protection activity via Fc-mediated effector functions against infections by diverse viruses, such as HIV^59,60^, SARS-CoV-2^61,62^, Zika virus^63^, Marburg virus^64^, West Nile virus^65^, and Crimean-Congo hemorrhagic fever virus^66^. For HA stem antibodies, our present study substantiates that the epitope location and approaching angle can influence Fc domain accessibility, hence the importance of Fc-mediated effector functions for protection activity *in vivo*. However, a recent study on HIV Env antibodies shows that their Fc-mediated effector functions are mainly determined by stoichiometry of binding and are not modulated by epitope location and approaching angle^67^. Whether the biophysical determinants for Fc-mediated effector functions are antigen-dependent warrants additional studies.

The recent shift in US funding policy against mRNA vaccine technology highlights the growing importance of broadly protective vaccine strategies for pandemic preparedness^68,69^. As exemplified during the COVID-19 pandemic, the mRNA vaccine technology has demonstrated an unprecedented ability to rapidly generate immunogens once an antigen sequence becomes available^70,71^. However, if mRNA vaccine technology becomes less accessible or deprioritized, pre-existing broad-spectrum immunity becomes more critical for pandemic preparedness. Vaccines that elicit antibodies targeting conserved epitopes, such as the influenza HA stem, offer a promising approach to mitigate the impact of unforeseeable influenza pandemic^72^. Therefore, the continued discovery and characterization of broadly protective influenza antibodies remain important, as they are key to the design of broadly protective influenza vaccines^72^.

## ACKNOWLEDGEMENTS

This work was supported by the Carl R. Woese Institute for Genomic Biology Postdoctoral Fellowship (H.L.), the Research Grants Council of the Hong Kong Special Administrative Region, China (HKU C7053-24G) (C.K.P.M), the Health and Medical Research Fund (no. 24230352) (C.K.P.M), the Vallee Scholars Program (N.C.W.), the Searle Scholars Program (N.C.W.), and Howard Hughes Medical Institute Emerging Pathogens Initiative (N.C.W.). We thank Kristen Flatt at the UIUC Materials Research Laboratory Central Research Facilities and Frank Vago at the Purdue Cryo-EM Facility for assistance with cryo-EM experiments.

## AUTHOR CONTRIBUTIONS

H.L., C.K.P.M., and N.C.W. conceived and designed the study. H.L., Y.W.H., T.P., Q.W.T., W.L., A.B.G., D.C., M.R.A., T.J.C.T., Y.S., A.M., M.S., J.J.H., E.X.M., L.E.W. and C.K.P.M. performed the experiments. H.L. C.K.P.M., and N.C.W. wrote the paper and all authors reviewed and/or edited the paper.

## METHODS

### Generation of phage display antibody libraries

Phage display antibody libraries were generated using a phagemid antibody library that we previously constructed from the peripheral blood mononuclear cells (PBMCs) of 245 healthy individuals^48^. Briefly, the phagemid antibody library was electroporated into E. coli TG1 cells, expanded, and infected with M13 KO7 ΔpIII hyperphage (hyperphage) or CM13 interference-resistant helper phage (CM phage) for phage packaging. The resulting phages were precipitated, purified, and resuspended in PBS to yield the phage display antibody libraries. The phage titers were determined by trypsin treatment, bacterial infection, and colony counting.

### Panning and screening of the phage display antibody libraries

100 ng of H1 mini-HA (in 100 µL of PBS) was added to each well of an Invitrogen Nunc MaxiSorp flat-bottom 96 well plate (Thermo Fisher Scientific), incubated overnight at 4°C. Hyperphage and CM phage sample was normalized to give approximately 10^11^ phage per 100 µL of reaction, which was then blocked with equal volume of 5% skim milk in PBS for 2 hours at room temperature. At the same time, wells in Maxisorp plate were blocked with 5% skim milk (added until the brim of wells) for 2 hours at room temperature. 200 µL of blocked phage sample was added to each well and incubated for 2 hours at 37°C. Wells were washed 5× with PBST (PBS supplemented with 0.1% Tween-20), 5× with PBS, tapped dry on paper towel after each wash. Phage were eluted from wells with 1 µg of TPCK-treated trypsin for 30 mins at 37°C. Eluted phage was added to 15 mL of E. coli TG1 culture outgrew to an OD_600_ of 0.45, followed by incubation at 37°C, standing for 30 mins, and shaking for 30 mins. Transductants were concentrated by centrifuging at 4500 ×g for 10 mins, and the cells as well as serial dilutions were platted onto 2YT agar supplemented with 100 µg mL^-1^ of ampicillin. Plates were incubated overnight at 30°C. Transductants were collected in 2YT medium supplemented with 30% glycerol and phage were prepared for 3 rounds of panning, as stated in the production of Fab-expressing phage. The stringency of screening was increased sequentially, by increasing the number of washes to 8× PBST + 7× PBS for the second round of panning, and 10× PBST + 10× PBS for the third round of panning.

### PacBio sequencing of the Fab libraries

The pre-selection Fab libraries and third round post-panning Fab libraries were amplified using PrimeSTAR Max DNA Polymerase (Takara Bio) per manufacturer’s instruction with the following primers (5’-GTA AAA CGA CGG CCA GTT TCA GTT TCG CTA CCG TGG CCC AAG CGG CC-3’ and 5’-CAG GAA ACA GCT ATG ACC CAC TAG TTT TTG TTC TTG GCC TGT TTG GCC ACA-3’). The PCR product was purified using a Monarch Gel Extraction Kit (New England Biolabs). A second round of PCR was carried out to add the sample index sequences to the amplicons (**Table S2)**. The final PCR products were sequenced on one SMRT cell 8M on a PacBio Revio system using the CCS sequencing mode and a 12-hour movie time.

### Analysis of PacBio sequencing data

Circular consensus sequences (CCSs) were generated from the raw subreads using SMRTLink v13.0, setting the parameters to require 99.9% accuracy and a minimum of 3 passes. CSSs in FASTQ format were parsed using the SeqIO module in BioPython and filtered based on the base calling quality score, where any read with more than five nucleotides of phred quality score <40 were removed. The adapter sequences were then identified on each read and trimmed from the Fab sequences. Reads that did not have the complete adapter sequences were also removed. The filtered reads were then aligned to the reference Fab sequences. Frequency (*F*) of Fab *i* of a given sample *s* was computed for each replicate as follows:

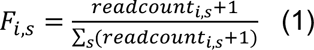

### Cell culture

HEK293T cells (human embryonic kidney cells, female) and MDCK-SIAT1 cells (Madin-Darby canine kidney cells with stable expression of human 2,6-sialtransferase, female, Sigma-Aldrich) were cultured in Dulbecco’s modified Eagle’s medium (DMEM) with high glucose (Thermo Fisher Scientific) supplemented with 10% heat-inactivated fetal bovine serum (FBS, Thermo Fisher Scientific), 1% penicillin-streptomycin (Thermo Fisher Scientific), and 1× GlutaMax (Thermo Fisher Scientific). Cell passaging was performed every 3 to 4 days using 0.05% Trypsin-ethylenediaminetetraacetic acid (EDTA) solution (Thermo Fisher Scientific). Expi293F cells (human embryonic kidney cells, female, Thermo Fisher Scientific) were maintained in Expi293 Expression Medium (Thermo Fisher Scientific). Sf9 cells (*Spodoptera frugiperda* ovarian cells, female, ATCC) were maintained in Sf-900 II SFM medium (Thermo Fisher Scientific).

### Expression and purification of mini-HA and HA

The H1 mini-HA (#4900)^34^, H3 mini-HA^49^, A/Solomon Island/3/2006 (H1N1) HA, A/Beijing/262/1995 (H1N1) HA, A/Michigan/45/2015 (H1N1) HA, A/Japan/305/1957 (H2N2) HA, and A/Texas/37/2024 (H5N1) HA were fused with N-terminal gp67 signal peptide and a C-terminal BirA biotinylation site, thrombin cleavage site, trimerization domain, and a 6× His-tag, and then cloned into a customized baculovirus transfer vector^73^. Subsequently, recombinant bacmid DNA was generated using the Bac-to-Bac system (Thermo Fisher Scientific) according to the manufacturer’s instructions. Baculovirus was generated by transfecting the purified bacmid DNA into adherent Sf9 cells using Cellfectin reagent (Thermo Fisher Scientific) according to the manufacturer’s instructions. The baculovirus was further amplified by passaging in adherent Sf9 cells at a multiplicity of infection (MOI) of 1. Recombinant mini-HA protein was expressed by infecting 1 L of suspension Sf9 cells at an MOI of 1. On day 3 post-infection, Sf9 cells were pelleted by centrifugation at 4000 ×*g* for 25 min, and soluble recombinant mini-HA and HA were purified from the supernatant by affinity chromatography using Ni Sepharose excel resin (Cytiva) and then size exclusion chromatography using a HiLoad 16/100 Superdex 200 prep grade column (Cytiva) in 20 mM Tris-HCl pH 8.0, 100 mM NaCl. The purified mini-HA protein was concentrated by Amicon spin filter (Millipore Sigma) and filtered by 0.22 µm centrifuge tube filters (Costar). Concentration of the protein was determined by nanodrop (Fisher Scientific). Proteins were subsequent aliquoted, flash frozen by dry-ice ethanol mixture, and stored at -80°C until used.

The following proteins were obtained from BEI Resources: A/California/07/2009 (H1N1) HA (NR-42635), A/bald eagle/Florida/W22–134-OP/2022 (H5N1) HA (NR-59476), A/northern pintail/WA/40964/2014 (H5N8) HA (NR-50174), and A/chicken/Netherlands/14015531/2014 (H5N8) HA (NR-50133). The following proteins were obtained from Sino Biological: A/USSR/90/1977 (H1N1) HA, A/Taiwan/01/1986 (H1N1), and A/Brisbane/02/2018 (H1N1) HA.

### Expression and purification of IgG and Fab

The heavy and light chain genes of the obtained antibody were synthesized as eBlocks (Integrated DNA Technologies), and then cloned into human IgG1 and human kappa or lambda light chain expression vectors using Gibson assembly according to a previously described method^74^. The plasmids of heavy chain and light chain were transiently co-transfected into Expi293F cells at a mass ratio of 2:1 (HC:LC) using ExpiFectamine 293 Reagent (Thermo Fisher Scientific). After transfection, the cell culture supernatant was collected at 6 days post-transfection. The IgG and Fab were then purified using a CaptureSelect CH1-XL pre-packed column (Thermo Fisher Scientific).

### Enzyme-linked immunosorbent assay (ELISA)

Nunc MaxiSorp ELISA plates (Thermo Fisher Scientific) were utilized and coated with 100 μL of recombinant proteins at a concentration of 1 μg ml^-1^ in a 1× PBS solution. The coating process was performed overnight at 4°C. On the following day, the ELISA plates were washed three times with 1× PBS supplemented with 0.05% Tween 20, and then blocked using 200 μL of 1× PBS with 5% non-fat milk powder for 2 hours at room temperature. After the blocking step, 100 μL of IgGs from the supernatant were added to each well and incubated for 2 hours at 37°C. The ELISA plates were washed three times to remove any unbound IgGs. Next, the ELISA plates were incubated with horseradish peroxidase (HRP)-conjugated goat anti-human IgG antibody (1:5000, Invitrogen) for 1 hour at 37°C. Subsequently, the ELISA plates were washed five times using PBS containing 0.05% Tween 20. Then, 100 μL of 1-Step Ultra TMB-ELISA Substrate Solution (Thermo Fisher Scientific) was added to each well. After 15 min incubation, 50 μL of 2 M H_2_SO_4_ solution was added to each well. The absorbance of each well was measured at a wavelength of 450 nm using a BioTek Synergy HTX Multimode Reader.

### Binding competition assay

Binding competition assays were performed by biolayer interferometry (BLI) using an Octet Red96e instrument (Sartorius) at room temperature, with modifications to a previously described method^75^. Briefly, His-tagged HA protein at 0.5 μM in 1× kinetics buffer (1× PBS, pH 7.4, 0.01% w/v BSA and 0.002% v/v Tween 20) was loaded onto anti-Penta-HIS (HIS1K) biosensors. The assay consisted of five steps: 1) baseline: 60 s with 1× kinetics buffer; 2) loading: 480 s with His-tagged HA protein; 3) baseline: 60 s with 1× kinetics buffer; 4) association: 120 s with antibody A in Fab format or with buffer; and 5) dissociation: 120 s with the antibody B in Fab format. For a given antibody B, its percentage competition with antibody A was computed as follow:

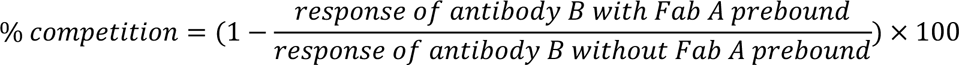

### Recombinant Virus Construction and Purification

Recombinant A/Puerto Rico/8/1934 (H1N1) (PR8) virus was rescued using the eight-plasmid influenza virus reverse genetics system^76^. Additionally, we rescued 7:1 reassortant viruses by using the HA segments from A/Chile/1/1983 (H1N1), A/Beijing/262/1995 (H1N1), A/Brisbane/59/2007 (H1N1), A/California/07/09 (H1N1), A/Michigan/45/2015 (H1N1), A/England/234600203/2023 (H1N2) and A/Vietnam/1203/2004 (H5N1) with the other seven segments from PR8 as previously described^77^. Briefly, DNA plasmids were transfected into a co-culture of HEK293T cells and MDCK-SIAT1 cells (ratio of 6:1) using Lipofectamine 2000 (Thermo Fisher Scientific) and incubated at 37°C for 48 hours. Recombinant viruses in the supernatant were plaque-purified on MDCK-SIAT1 cells. Individual plaques were picked and incubated with fresh MDCK-SIAT1 cells, and viral RNAs were extracted from supernatant at 72 hours post-infection. The sequences of HA segments were confirmed by Sanger sequencing. All experiments involving H5 viruses were performed in the Biosafety Level 3 (BSL3) facility at The Chinese University of Hong Kong.

### Virus neutralization assay

MDCK-SIAT1 cells were seeded in a 96-well, flat-bottom cell culture plate (Thermo Fisher Scientific). The next day, serially diluted monoclonal antibodies were mixed with an equal volume of virus and incubated at 37°C for 1 hour. The antibody-virus mixture was then incubated with MDCK-SIAT1 cells at 37°C for 1 hour after the cells were washed twice with PBS. Subsequently, the antibody-virus mixture was replaced with Minimum Essential Medium (MEM) supplemented with 25 mM of 4-(2-hydroxyethyl)-1-piperazineethanesulfonic acid (HEPES) and 1 μg mL^-1^ of Tosyl phenylalanyl chloromethyl ketone (TPCK)-trypsin. The plate was incubated at 37°C for 72 hours and the presence of virus was detected by hemagglutination assay. The results were analyzed using Prism software (GraphPad).

### ADCC reporter bioassay

1.5 × 10^4^ MDCK cells were seeded in white, flat bottom 96-well cell culture plates (Thermo Fisher) and incubated at 37°C overnight. The next day, cells were washed three times with PBS and 100 μL of PR8 virus at 1.5 × 10^6^ plaque forming units (PFU) mL^-1^ in MEM was added to each well. After incubating the plate for 24 hours at 37°C, the media was removed and 25 μL of serially diluted antibodies in 1:10 in Roswell Park Memorial Institute (RPMI) 1640 media (Thermo Fisher), 25 μL of effector cells (engineered Jurkat cells stably expressing human FcγRIIIa V158 and NFAT-induced luciferase) and 25 μL of RPMI 1640 media were added. After 6 hours incubation at 37°C, 75 μL of Bio-Glo luciferase (Promega) was added to each well. The plate was incubated for 10 min in the dark and the luciferase induced luminescence measured with a BioTek Synergy HTX Multimode Reader (Agilent). Data were analyzed using Prism software and the area under the curve (AUC) values were determined.

### Prophylactic and therapeutic protection experiments

Female BALB/c mice at 6 weeks old (n = 5 mice/group) were anesthetized with isoflurane and intranasally infected with 5× median lethal dose (LD_50_) of PR8 H1N1 virus or PR8-H5N1 (7:1 on backbone from A/Puerto Rico/8/1934). Mice were given the indicated antibody at a dose of 5 mg kg^-1^ intraperitoneally at 4 hours before infection (prophylaxis) or 24 hours after infection (therapeutics). Weight loss was monitored daily for 14 days. The humane endpoint was defined as a weight loss of 25% from initial weight on day 0. Of note, while our BALB/c mice were not modified to facilitate interaction with human IgG1, human IgG1 could interact with mouse Fc gamma receptors^78,79^. To determine the lung viral titers on day 3 post-infection, lungs of infected mice were harvested and homogenized in 1 mL of MEM with 1 μg mL^-1^ of TPCK-trypsin with gentleMACS Dissociator (Miltenyi Biotec). Subsequently, viral titers were measured by TCID_50_ (median tissue culture infectious dose) assay. The results were analyzed using Prism software. The animal experiments were performed in accordance with protocols approved by UIUC Institutional Animal Care and Use Committee (IACUC). All mouse experiments involving H5 viruses were conducted in the Biosafety Level 3 (BSL3) facility at The Chinese University of Hong Kong.

### Cryo-EM sample preparation and data collection

The purified A/Solomon Islands/3/2006 (H1N1) HA protein was mixed with the Fab at 1:4 molar ratio (one HA trimer per four Fabs) and incubated overnight at 4°C before purifying by size exclusion chromatography on the Superose 6 Increased 10/300 column (Cytiva) in 20 mM Tris-HCl pH 8.0 and 100 mM NaCl. For the complexes with HB31 and HB34, anchor Fab FISW84^53^ was also added to reduce the preferred orientation. The complex peak was concentrated to ∼3 mg mL^-1^ and mixed with 0.5% w/v n-octyl-ß-D-glucoside (Anagrade) to a final concentration of 0.1% w/v just before loading on the grid. Cryo-EM grids were prepared using a Vitrobot Mark IV machine. An aliquot of 3 µL sample was applied to a 300-mesh Quantifoil R1.2/1.3 Cu grid pretreated with glow-discharge. High-resolution cryo-EM movies were collected on an FEI Titan Krios at 300 kV with a Gatan K3 detector.

### Cryo-EM image processing and model building

Data processing was performed with CryoSPARC (version 4.5)^80^. Movies were subjected to motion correction and CTF estimation, and particles were picked with CryoSPARC blob picker followed by 2D classification. Best classes from blob picker were used as templates for CryoSPARC template pickers, and the resulting particles were cleaned up by multiple rounds of 2D classification before ab initio reconstruction. The best class from ab initio reconstruction was subjected to homogenous refinement, reference-based motion correction, another round of homogenous refinement, local and global CTF estimation, and non-uniform refinement. All datasets except for HB31 were processed with C3 symmetry. Due to substoichiometric quantities of bound FISW84 Fab in the HB31 complex, the data were processed in C1 symmetry, and FISW84 Fab was not built into the map. All maps were sharpened with DeepEMhancer^81^, and all initial atomic model were built using ModelAngelo^82^. The models were subjected to multiple rounds of manual refinement in Coot (version 0.9.8)^83^ and real-space refinement in Phenix^84^. This process was iterated for several cycles until no significant improvement of the model was observed.

### Biolayer interferometry binding assay

Binding assays were performed by biolayer interferometry (BLI) using an Octet Red96e instrument (Sartorius) at room temperature as described previously. Briefly, the indicated Fabs at 300 nM, 100 nM, and 33 nM in 1× kinetics buffer (PBS at pH 7.4, 0.01% w/v BSA, and 0.002% v/v Tween 20) were loaded onto Anti-Human Fab-CH1 2nd Generation (FAB2G) biosensors and incubated with A/Texas/37/2024 (H5N1) HA and H1 stem at 20 µg mL^-1^. The assay consisted of five steps, each run for at least 60 seconds: (1) baseline: 1× kinetics buffer; (2) loading: Fab proteins; (3) baseline: 1× kinetics buffer; (4) association: HA proteins; and (5) dissociation: 1× kinetics buffer. For estimating the K_D_, a 1:1 binding model was used.

## DATA AVAILABILITY

PacBio sequencing data have been submitted to the NIH Short Read Archive under accession number: BioProject PRJNA1104662. Cryo-EM maps have been deposited in the Electron Microscopy Data Bank with accession numbers: EMD-70809, EMD-70810, and EMD-70811. The refined models have been deposited in the Protein Data Bank with accession numbers: 9OSS, 9OST, and 9OSU. Custom scripts for analyzing the PacBio sequencing data have been deposited to https://github.com/Karobben/miniHA_Abs_Phage_Screen.

## REFERENCES

1 Origanization, W. H. The burden of influenza. https://www.who.int/news-room/feature-stories/detail/the-burden-of-influenza (2024).

2 Barr, I. G. & Subbarao, K. Implications of the apparent extinction of B/Yamagata-lineage human influenza viruses. NPJ Vaccines 9, 219, doi:10.1038/s41541-024-01010-y (2024).

3 Caini, S. et al. Probable extinction of influenza B/Yamagata and its public health implications: a systematic literature review and assessment of global surveillance databases. Lancet Microbe 5, 100851, doi:10.1016/S2666-5247(24)00066-1 (2024).

4 Saunders-Hastings, P. R. & Krewski, D. Reviewing the history of pandemic influenza: understanding patterns of emergence and transmission. Pathogens 5, doi:10.3390/pathogens5040066 (2016).

5 Writing Committee of the Second World Health Organization Consultation on Clinical Aspects of Human Infection with Avian Influenza, A. V. et al. Update on avian influenza A (H5N1) virus infection in humans. N Engl J Med 358, 261–273, doi:10.1056/NEJMra0707279 (2008).

6 Burrough, E. R. et al. Highly pathogenic avian influenza A (H5N1) clade 2.3.4.4b virus infection in domestic dairy cattle and cats, United States, 2024. Emerg Infect Dis 30, 1335–1343, doi:10.3201/eid3007.240508 (2024).

7 Basilio, H. Bird-flu vaccine for cattle aces early test. Nature, doi:10.1038/d41586-025-01497-y (2025).

8 CDC. H5 bird flu: Current situation. https://www.cdc.gov/bird-flu/situation-summary/index.html (2025).

9 Krammer, F. The human antibody response to influenza A virus infection and vaccination. Nat Rev Immunol 19, 383–397, doi:10.1038/s41577-019-0143-6 (2019).

10 Wu, N. C. & Wilson, I. A. Influenza hemagglutinin structures and antibody recognition. Cold Spring Harb Perspect Med 10, doi:10.1101/cshperspect.a038778 (2020).

11 Wu, N. C. & Wilson, I. A. A perspective on the structural and functional constraints for immune evasion: insights from influenza virus. J Mol Biol 429, 2694–2709, doi:10.1016/j.jmb.2017.06.015 (2017).

12 Roubidoux, E. K. et al. Mutations in the hemagglutinin stalk domain do not permit escape from a protective, stalk-based vaccine-induced immune response in the mouse model. mBio 12, doi:10.1128/mBio.03617-20 (2021).

13 Doud, M. B., Lee, J. M. & Bloom, J. D. How single mutations affect viral escape from broad and narrow antibodies to H1 influenza hemagglutinin. Nat Commun 9, 1386, doi:10.1038/s41467-018-03665-3 (2018).

14 Dreyfus, C. et al. Highly conserved protective epitopes on influenza B viruses. Science 337, 1343–1348, doi:10.1126/science.1222908 (2012).

15 Kallewaard, N. L. et al. Structure and function analysis of an antibody recognizing all influenza A subtypes. Cell 166, 596–608, doi:10.1016/j.cell.2016.05.073 (2016).

16 Corti, D. et al. A neutralizing antibody selected from plasma cells that binds to group 1 and group 2 influenza A hemagglutinins. Science 333, 850–856, doi:10.1126/science.1205669 (2011).

17 Andrews, S. F. et al. Preferential induction of cross-group influenza A hemagglutinin stem-specific memory B cells after H7N9 immunization in humans. Sci Immunol 2, doi:10.1126/sciimmunol.aan2676 (2017).

18 Ekiert, D. C. et al. A highly conserved neutralizing epitope on group 2 influenza A viruses. Science 333, 843–850, doi:10.1126/science.1204839 (2011).

19 Friesen, R. H. et al. A common solution to group 2 influenza virus neutralization. Proc Natl Acad Sci U S A 111, 445–450, doi:10.1073/pnas.1319058110 (2014).

20 Lang, S. et al. Antibody 27F3 broadly targets influenza A group 1 and 2 hemagglutinins through a further variation in V(H)1-69 antibody orientation on the HA stem. Cell Rep 20, 2935–2943, doi:10.1016/j.celrep.2017.08.084 (2017).

21 Joyce, M. G. et al. Vaccine-induced antibodies that neutralize group 1 and group 2 influenza A viruses. Cell 166, 609–623, doi:10.1016/j.cell.2016.06.043 (2016).

22 Wu, N. C. et al. Convergent evolution in breadth of two V(H)6-1-encoded influenza antibody clonotypes from a single donor. Cell Host Microbe 28, 434–444 e434, doi:10.1016/j.chom.2020.06.003 (2020).

23 Chuang, G. Y. et al. Sequence-signature optimization enables improved identification of human HV6-1-derived class antibodies that neutralize diverse influenza A viruses. Front Immunol 12, 662909, doi:10.3389/fimmu.2021.662909 (2021).

24 Wu, N. C. et al. Recurring and adaptable binding motifs in broadly neutralizing antibodies to influenza virus are encoded on the D3-9 segment of the Ig gene. Cell Host Microbe 24, 569–578 e564, doi:10.1016/j.chom.2018.09.010 (2018).

25 Wang, Y. et al. An explainable language model for antibody specificity prediction using curated influenza hemagglutinin antibodies. Immunity 57, 2453–2465 e2457, doi:10.1016/j.immuni.2024.07.022 (2024).

26 Ekiert, D. C. et al. Antibody recognition of a highly conserved influenza virus epitope. Science 324, 246–251, doi:10.1126/science.1171491 (2009).

27 Andrews, S. F. et al. A single residue in influenza virus H2 hemagglutinin enhances the breadth of the B cell response elicited by H2 vaccination. Nat Med 28, 373–382, doi:10.1038/s41591-021-01636-8 (2022).

28 Sangesland, M. et al. Allelic polymorphism controls autoreactivity and vaccine elicitation of human broadly neutralizing antibodies against influenza virus. Immunity 55, 1693–1709 e1698, doi:10.1016/j.immuni.2022.07.006 (2022).

29 Guthmiller, J. J. et al. Broadly neutralizing antibodies target a haemagglutinin anchor epitope. Nature 602, 314–320, doi:10.1038/s41586-021-04356-8 (2022).

30 Lin, T. H. et al. Structurally convergent antibodies derived from different vaccine strategies target the influenza virus HA anchor epitope with a subset of V(H)3 and V(K)3 genes. Nat Commun 16, 1268, doi:10.1038/s41467-025-56496-4 (2025).

31 Kadam, R. U. et al. Potent peptidic fusion inhibitors of influenza virus. Science 358, 496–502, doi:10.1126/science.aan0516 (2017).

32 Krammer, F., Pica, N., Hai, R., Margine, I. & Palese, P. Chimeric hemagglutinin influenza virus vaccine constructs elicit broadly protective stalk-specific antibodies. J Virol 87, 6542–6550, doi:10.1128/JVI.00641-13 (2013).

33 Yassine, H. M. et al. Hemagglutinin-stem nanoparticles generate heterosubtypic influenza protection. Nat Med 21, 1065–1070, doi:10.1038/nm.3927 (2015).

34 Impagliazzo, A. et al. A stable trimeric influenza hemagglutinin stem as a broadly protective immunogen. Science 349, 1301–1306, doi:10.1126/science.aac7263 (2015).

35 Widge, A. T. et al. An influenza hemagglutinin stem nanoparticle vaccine induces cross-group 1 neutralizing antibodies in healthy adults. Sci Transl Med 15, eade4790, doi:10.1126/scitranslmed.ade4790 (2023).

36 Andrews, S. F. et al. An influenza H1 hemagglutinin stem-only immunogen elicits a broadly cross-reactive B cell response in humans. Sci Transl Med 15, eade4976, doi:10.1126/scitranslmed.ade4976 (2023).

37 Nachbagauer, R. et al. A chimeric hemagglutinin-based universal influenza virus vaccine approach induces broad and long-lasting immunity in a randomized, placebo-controlled phase I trial. Nat Med 27, 106–114, doi:10.1038/s41591-020-1118-7 (2021).

38 Mantus, G. E. et al. Vaccination with different group 2 influenza subtypes alters epitope targeting and breadth of hemagglutinin stem-specific human B cells. Sci Transl Med 17, eadr8373, doi:10.1126/scitranslmed.adr8373 (2025).

39 Fleishman, S. J. et al. Computational design of proteins targeting the conserved stem region of influenza hemagglutinin. Science 332, 816–821, doi:10.1126/science.1202617 (2011).

40 Whitehead, T. A. et al. Optimization of affinity, specificity and function of designed influenza inhibitors using deep sequencing. Nat Biotechnol 30, 543–548, doi:10.1038/nbt.2214 (2012).

41 Chevalier, A. et al. Massively parallel de novo protein design for targeted therapeutics. Nature 550, 74–79, doi:10.1038/nature23912 (2017).

42 Koday, M. T. et al. A computationally designed hemagglutinin stem-binding protein provides in vivo protection from influenza independent of a host immune response. PLoS Pathog 12, e1005409, doi:10.1371/journal.ppat.1005409 (2016).

43 Bennett, N. R. et al. Atomically accurate de novo design of single-domain antibodies. bioRxiv, 2024.2003.2014.585103, doi:10.1101/2024.03.14.585103 (2024).

44 Pascha, M. N. et al. Inhibition of H1 and H5 influenza A virus entry by diverse macrocyclic peptides targeting the hemagglutinin stem region. ACS Chem Biol 17, 2425–2436, doi:10.1021/acschembio.2c00040 (2022).

45 Kitamura, S. et al. Ultrapotent influenza hemagglutinin fusion inhibitors developed through SuFEx-enabled high-throughput medicinal chemistry. Proc Natl Acad Sci U S A 121, e2310677121, doi:10.1073/pnas.2310677121 (2024).

46 van Dongen, M. J. P. et al. A small-molecule fusion inhibitor of influenza virus is orally active in mice. Science 363, doi:10.1126/science.aar6221 (2019).

47 Yao, Y. et al. An influenza A hemagglutinin small-molecule fusion inhibitor identified by a new high-throughput fluorescence polarization screen. Proc Natl Acad Sci U S A 117, 18431–18438, doi:10.1073/pnas.2006893117 (2020).

48 Lv, H. et al. Evolution of antibody cross-reactivity to influenza H5N1 neuraminidase from an N2-specific germline. Cell Host Microbe, 10.1016/j.chom.2025.09.015 (2025).

49 Corbett, K. S. et al. Design of nanoparticulate group 2 influenza virus hemagglutinin stem antigens that activate unmutated ancestor B cell receptors of broadly neutralizing antibody lineages. mBio 10, doi:10.1128/mBio.02810-18 (2019).

50 Henry Dunand, C. J., et al. Both neutralizing and non-neutralizing human H7N9 influenza vaccine-induced monoclonal antibodies confer protection. Cell Host Microbe 19, 800–813, doi:10.1016/j.chom.2016.05.014 (2016).

51 Sutton, T. C. et al. In vitro neutralization is not predictive of prophylactic efficacy of broadly neutralizing monoclonal antibodies CR6261 and CR9114 against lethal H2 influenza virus challenge in mice. J Virol 91, doi:10.1128/JVI.01603-17 (2017).

52 Lo, M. et al. Effector-attenuating substitutions that maintain antibody stability and reduce toxicity in mice. J Biol Chem 292, 3900–3908, doi:10.1074/jbc.M116.767749 (2017).

53 Benton, D. J. et al. Influenza hemagglutinin membrane anchor. Proc Natl Acad Sci U S A 115, 10112–10117, doi:10.1073/pnas.1810927115 (2018).

54 McIntire, K. M. et al. Maturation of germinal center B cells after influenza virus vaccination in humans. J Exp Med 221, doi:ARTN e20240668 10.1084/jem.20240668 (2024).

55 Ataca, S. et al. Modulating the immunodominance hierarchy of immunoglobulin germline-encoded structural motifs targeting the influenza hemagglutinin stem. Cell Rep 43, 114990, doi:10.1016/j.celrep.2024.114990 (2024).

56 Yuan, M. & Wilson, I. A. The D gene in CDR H3 determines a public class of human antibodies to SARS-CoV-2. Vaccines (Basel) 12, doi:10.3390/vaccines12050467 (2024).

57 Wang, Y. et al. A large-scale systematic survey reveals recurring molecular features of public antibody responses to SARS-CoV-2. Immunity 55, 1105–1117 e1104, doi:10.1016/j.immuni.2022.03.019 (2022).

58 Carter, J. J. et al. Human monoclonal antibodies to HPV16 show evidence for common developmental pathways and public epitopes. PLoS Pathog 21, e1013086, doi:10.1371/journal.ppat.1013086 (2025).

59 Horwitz, J. A. et al. Non-neutralizing antibodies alter the course of HIV-1 infection in vivo. Cell 170, 637–648 e610, doi:10.1016/j.cell.2017.06.048 (2017).

60 Mayr, L. M., Su, B. & Moog, C. Non-neutralizing antibodies directed against HIV and their functions. Front Immunol 8, 1590, doi:10.3389/fimmu.2017.01590 (2017).

61 Pierre, C. N. et al. Non-neutralizing SARS-CoV-2 N-terminal domain antibodies protect mice against severe disease using Fc-mediated effector functions. PLoS Pathog 20, e1011569, doi:10.1371/journal.ppat.1011569 (2024).

62 Clark, J. J. et al. Protective effect and molecular mechanisms of human non-neutralizing cross-reactive spike antibodies elicited by SARS-CoV-2 mRNA vaccination. Cell Rep 43, 114922, doi:10.1016/j.celrep.2024.114922 (2024).

63 Bailey, M. J. et al. Human antibodies targeting Zika virus NS1 provide protection against disease in a mouse model. Nat Commun 9, 4560, doi:10.1038/s41467-018-07008-0 (2018).

64 Ilinykh, P. A. et al. Non-neutralizing Antibodies from a Marburg Infection Survivor Mediate Protection by Fc-Effector Functions and by Enhancing Efficacy of Other Antibodies. Cell Host Microbe 27, 976–991 e911, doi:10.1016/j.chom.2020.03.025 (2020).

65 Chung, K. M. et al. Antibodies against West Nile Virus nonstructural protein NS1 prevent lethal infection through Fc gamma receptor-dependent and -independent mechanisms. J Virol 80, 1340–1351, doi:10.1128/JVI.80.3.1340-1351.2006 (2006).

66 Golden, J. W. et al. GP38-targeting monoclonal antibodies protect adult mice against lethal Crimean-Congo hemorrhagic fever virus infection. Sci Adv 5, eaaw9535, doi:10.1126/sciadv.aaw9535 (2019).

67 Bick, M. V. et al. Molecular parameters governing antibody FcgammaR signaling and effector functions in the context of HIV envelope. Cell Rep 44, 115331, doi:10.1016/j.celrep.2025.115331 (2025).

68 US government slams mRNA vaccine research. Nat Biotechnol 43, 1400, doi:10.1038/s41587-025-02826-2 (2025).

69 Petersen, E. et al. Protecting the future of vaccine development amidst US funding withdrawal for mRNA vaccine research. Lancet Microbe, 101226, doi:10.1016/j.lanmic.2025.101226 (2025).

70 Corbett, K. S. et al. SARS-CoV-2 mRNA vaccine design enabled by prototype pathogen preparedness. Nature 586, 567–571, doi:10.1038/s41586-020-2622-0 (2020).

71 Polack, F. P. et al. Safety and efficacy of the BNT162b2 mRNA Covid-19 vaccine. N Engl J Med 383, 2603–2615, doi:10.1056/NEJMoa2034577 (2020).

72 Wu, N. C. & Wilson, I. A. Structural insights into the design of novel anti-influenza therapies. Nat Struct Mol Biol 25, 115–121, doi:10.1038/s41594-018-0025-9 (2018).

73 Ekiert, D. C. et al. A highly conserved neutralizing epitope on group 2 influenza A viruses. Science 333, 843–850, doi:10.1126/science.1204839 (2011).

74 Guthmiller, J. J., Dugan, H. L., Neu, K. E., Lan, L. Y. & Wilson, P. C. An efficient method to generate monoclonal antibodies from human B cells. Methods Mol Biol 1904, 109–145, doi:10.1007/978-1-4939-8958-4_5 (2019).

75 Teo, Q. W. et al. Stringent and complex sequence constraints of an IGHV1-69 broadly neutralizing antibody to influenza HA stem. Cell Rep 42, 113410, doi:10.1016/j.celrep.2023.113410 (2023).

76 Hoffmann, E., Neumann, G., Kawaoka, Y., Hobom, G. & Webster, R. G. A DNA transfection system for generation of influenza A virus from eight plasmids. Proc Natl Acad Sci U S A 97, 6108–6113, doi:10.1073/pnas.100133697 (2000).

77 Lv, H. et al. Differential antigenic imprinting effects between influenza H1N1 hemagglutinin and neuraminidase in a mouse model. J Virol 99, e0169524, doi:10.1128/jvi.01695-24 (2025).

78 Overdijk, M. B. et al. Crosstalk between human IgG isotypes and murine effector cells. J Immunol 189, 3430–3438, doi:10.4049/jimmunol.1200356 (2012).

79 Dekkers, G. et al. Affinity of human IgG subclasses to mouse Fc gamma receptors. MAbs 9, 767–773, doi:10.1080/19420862.2017.1323159 (2017).

80 Punjani, A., Rubinstein, J. L., Fleet, D. J. & Brubaker, M. A. cryoSPARC: algorithms for rapid unsupervised cryo-EM structure determination. Nat Methods 14, 290–296, doi:10.1038/nmeth.4169 (2017).

81 Sanchez-Garcia, R. et al. DeepEMhancer: a deep learning solution for cryo-EM volume post-processing. Commun Biol 4, 874, doi:10.1038/s42003-021-02399-1 (2021).

82 Jamali, K. et al. Automated model building and protein identification in cryo-EM maps. Nature 628, 450–457, doi:10.1038/s41586-024-07215-4 (2024).

83 Casanal, A., Lohkamp, B. & Emsley, P. Current developments in Coot for macromolecular model building of Electron Cryo-microscopy and Crystallographic Data. Protein Sci 29, 1069–1078, doi:10.1002/pro.3791 (2020).

84 Afonine, P. V. et al. Real-space refinement in PHENIX for cryo-EM and crystallography. Acta Crystallogr D Struct Biol 74, 531–544, doi:10.1107/S2059798318006551 (2018).

