## Supplementary material for "Structure basis for distinct protective mechanisms of IGHV3-23 antibodies targeting influenza hemagglutinin stem": SI figure

**Table S1. Cryo-EM data collection, refinement and validation statistics.**

| | HB31 Fab + SI06 HA + FISW84 Fab | HB34 Fab + SI06 HA + FISW84 Fab | HB315 Fab + SI06 HA |
|---|---|---|---|
| **Data collection and processing** | | | |
| Magnification | 81,000 | 81,000 | 81,000 |
| Voltage (kV) | 300 | 300 | 300 |
| Electron exposure ($e^{-}$/Å$^{2}$) | 57.35 | 57.35 | 57.35 |
| Defocus range (μm) | -0.5 – 3.0 | -0.5 – 3.0 | -0.5 – 3.0 |
| Pixel size (Å) | 0.529 | 0.529 | 0.529 |
| Symmetry imposed | C1 | C3 | C3 |
| Initial particle images (no.) | 599,816 | 714,635 | 647,100 |
| Final particle images (no.) | 406,141 | 304,985 | 300,983 |
| Map resolution (Å) | 2.76 | 2.64 | 2.67 |
| FSC threshold 0.143 | | | |
| Map postprocessing | DeepEMhancer | DeepEMhancer | DeepEMhancer |
| **Refinement** | | | |
| Initial model used (PDB code) | N/A | N/A | N/A |
| Model composition | | | |
| Non-hydrogen atoms | 16,352 | 20,816 | 16,181 |
| Protein residues | 2,161 | 2,753 | 2,154 |
| Root-mean-square deviations | | | |
| Bond lengths (Å) | 0.003 | 0.003 | 0.004 |
| Bond angles (°) | 0.526 | 0.547 | 0.581 |
| Validation | | | |
| MolProbity score | 1.95 | 2.05 | 2.15 |
| Clashscore | 7.19 | 6.34 | 5.89 |
| Poor rotamers (%) | 2.04 | 3.00 | 4.45 |
| Ramachadran plot | | | |
| Favored (%) | 95.35 | 95.17 | 95.12 |
| Allowed (%) | 4.51 | 4.83 | 4.84 |
| Disallowed (%) | 0.14 | 0.00 | 0.05 |
| **PDB code** | 9OSS | 9OST | 9OSU |
| **EMDB code** | EMD-70809 | EMD-70810 | EMD-70811 |

**Table S2. Sequences of sample indexing primers.**

| Primer name | Sequence (5’ to 3’) |
|---|---|
| PacBio-F-1 | TGAACCTTGTAAAACGACGGCCAGTTTCAG |
| PacBio-F-2 | TGCTAAGTGTAAAACGACGGCCAGTTTCAG |
| PacBio-F-3 | TGTTCTCTGTAAAACGACGGCCAGTTTCAG |
| PacBio-R-1 | GTCGTGATCAGGAAACAGCTATGACCCACT |
| PacBio-R-2 | ACCACTGTCAGGAAACAGCTATGACCCACT |
| PacBio-R-3 | TGGATCTGCAGGAAACAGCTATGACCCACT |

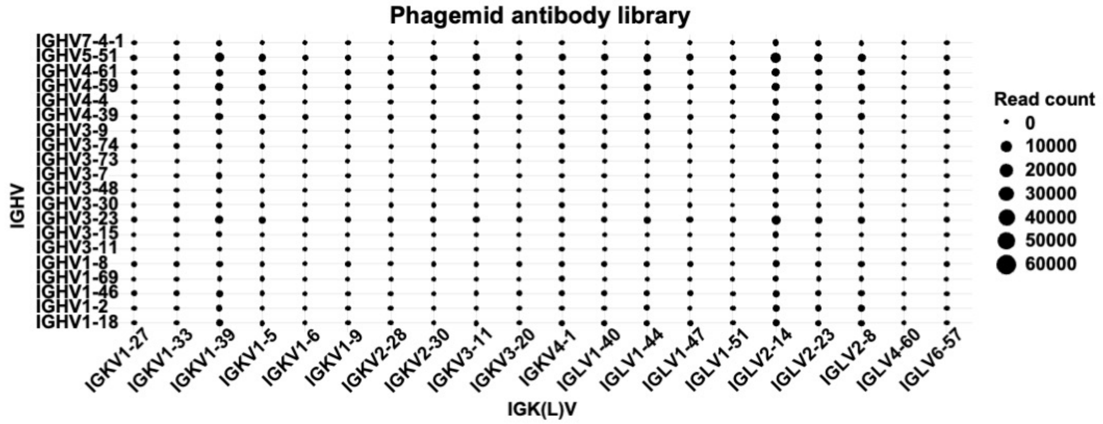


**Figure S1. Distribution of IGHV/IGK(L)V pairings in the phagemid antibody library.** Point size reflects the sequencing read count for each IGHV/IGK(L)V combination.

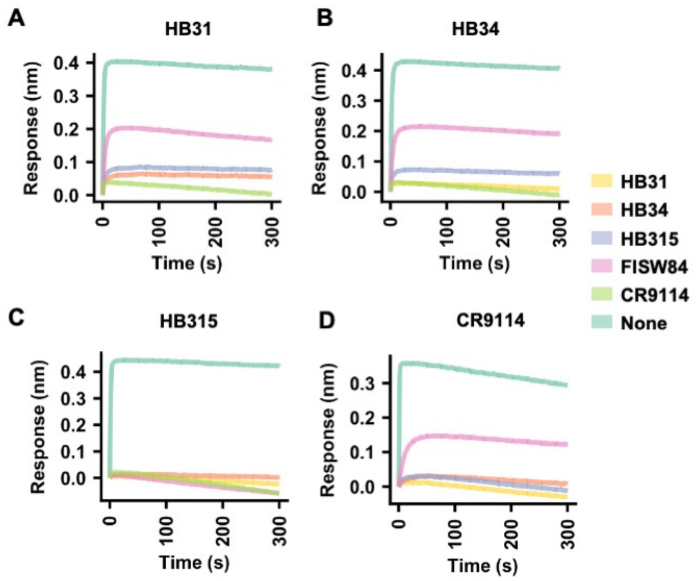


**Figure S2. Sensorgrams of the antibody competition assay.** Competition between antibodies was determined by biolayer interferometry. Briefly, A/Michigan/45/2015 HA was immobilized on the biosensor, followed by the binding of **(A)** HB31, **(B)** HB34, **(C)** HB315, and **(D)** CR9114. Subsequently, the indicated antibodies as color coded were added to test for competition. Of note, CR9114 and FISW84 bind to the central stem epitope[1] and anchor epitope[2], respectively. Sensorgram of the association step of the second antibody. All antibodies were in Fab format.

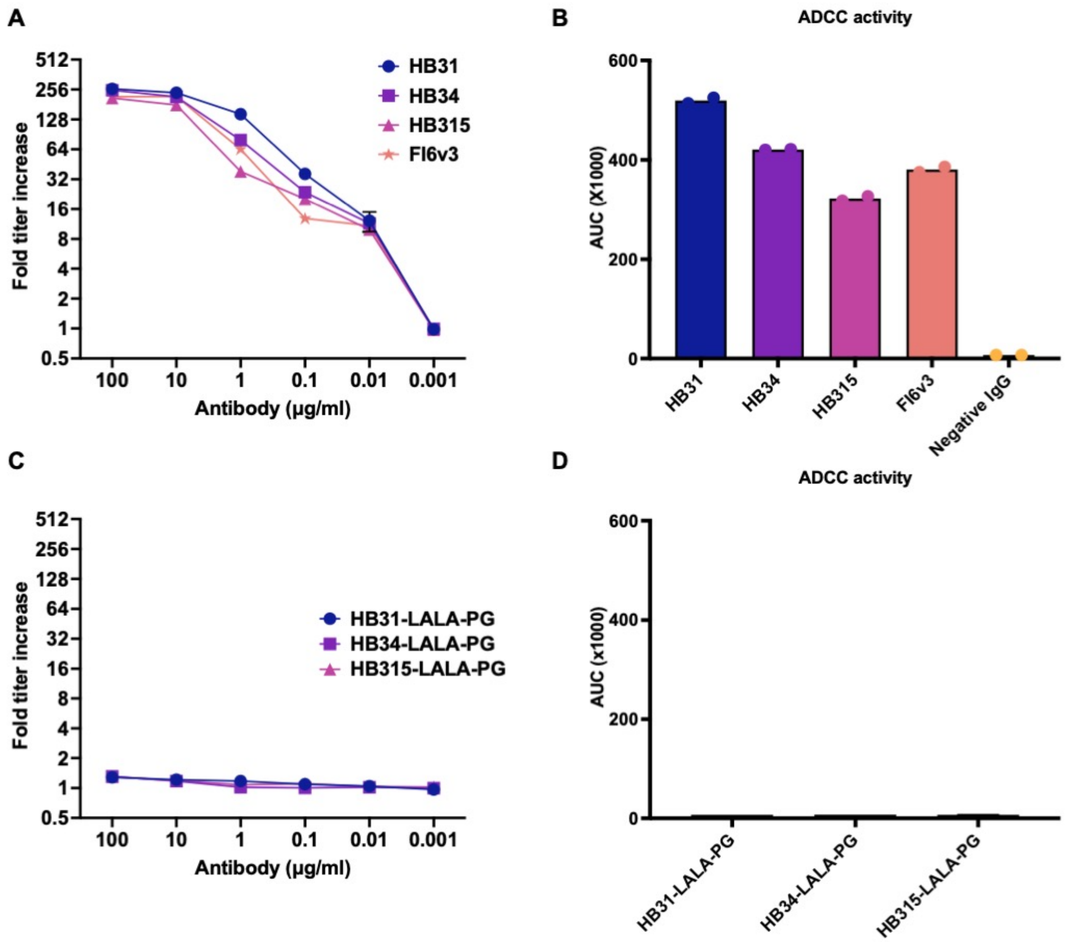


**Figure S3. Antibody-dependent cellular cytotoxicity (ADCC) report assay.** ADCC activity of the indicated antibodies was measured. FI6v3[3], which is a HA stem antibody known to have ADCC activity, was used as a positive control. IgG from human serum was used as a negative control. **(A and C)** Titration curve and **(B and D)** area under the curve (AUC) are shown.

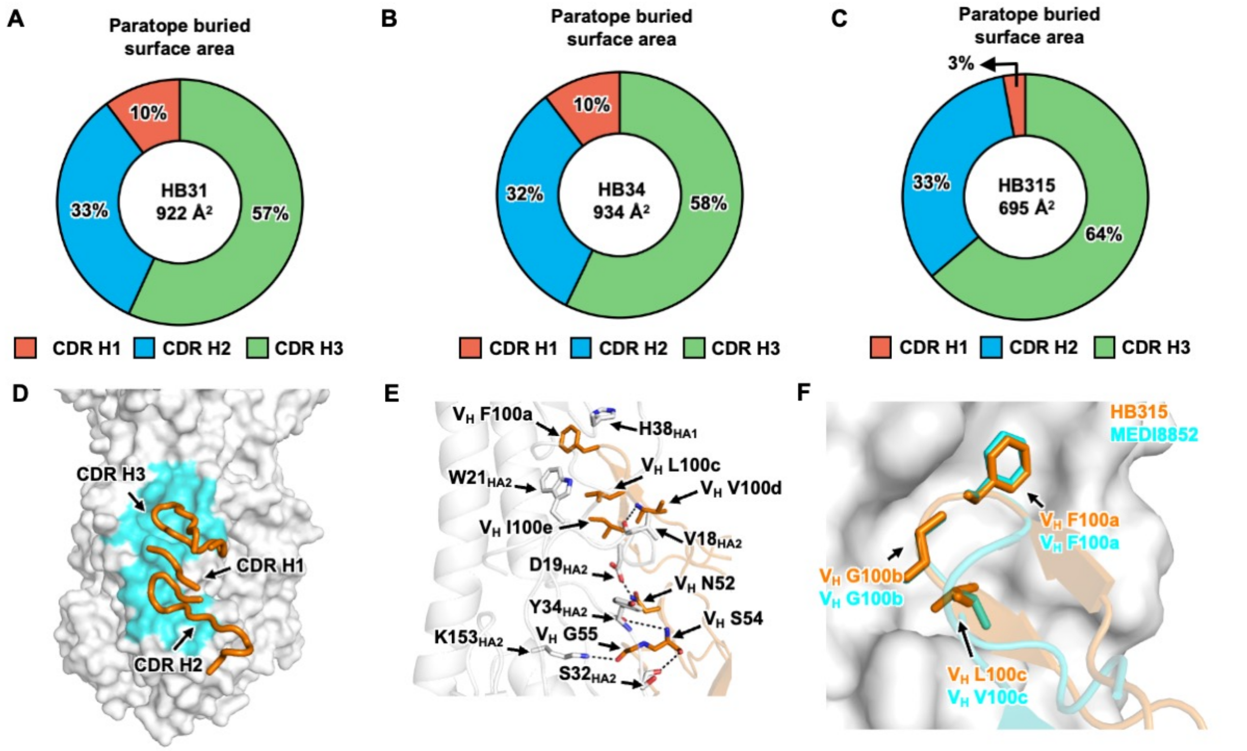


**Figure S4. Structural analysis of HB31, HB34, and HB315.** Contributions of different CDRs of **(A)** HB31, **(B)** HB34, and **(C)** HB315 to the paratope buried surface area are shown. **(D)** HA is shown as surface representation with the HB315 epitope colored in cyan. The heavy chain CDRs of HB315 are shown as orange cartoon. **(E)** Key residues at the interface between HA and HB315 are shown. H-bonds are shown black dashed lines. **(F)** Interactions between HA (white surface) and the FG[L/V] motifs in the CDR H3s of MEDI8852 (PDB 5JW4, cyan)[4] and HB315 (orange) are shown.

**A**

**HB31/HB34**

```
                     C  A  R  D  R  P  V  L  R  Y  F  D  W  Q  P  Y  G  L  D  V  W
              HB31:TGTGCGAGAGATCGCCCCGTGTTACGATATTTTGACTGGCAACCATACGGTTTGGACGTCTGG
Germline sequence:TGTGCGAAA           GTGTTACGATATTTTGACTGG     TACGGTTTGGACGTCTGG
                  IGHV3                     IGHD3-9                   IGHJ6
```

**B**

**HB315**

```
                     C  A  K  D  L  G  R  A  I  F  G  L  V  I  P  E  G  A  F  D  I  W
              HB31:TGTGCGAAAGACTTAGGGCGGGCGATTTTTGGACTGGTTATCCCAGAGGGTGCTTTTGATATCTGG
Germline sequence:TGTGCGAAAGA           CGATTTTTGGACTGGTTAT           TGCTTTTGATATCTGG
                  IGHV3                      IGHD3-3                       IGHJ3
```

**Figure S5. Putative germline sequences for the CDR H3s of** HB31, HB34, and HB315**.** Amino acid and nucleotide sequences of the heavy chain V-D-J junction are shown for **(A)** HB31, HB34, and **(C)** HB315. Putative germline sequences and segments for IGHV, IGHD, and IGHJ, are indicated and colored in blue, purple, and red, respectively. Somatically mutated nucleotides are underlined. Intervening spaces at the V-D and D-J junctions are N-nucleotide additions.

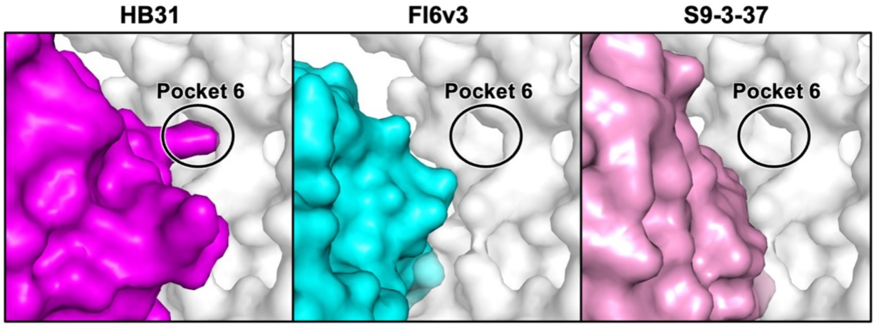


**Figure S6. Structure comparison of *IGHD3-9* HA stem antibodies.** *IGHD3-9* HA stem antibodies HB31 (magenta), FI6v3 (PDB 3ZTN, cyan)[3] and S9-3-37 (PDB 6E3H, pink)[5], as well as HA (white) are shown as surface representation. The location of pocket 6 is indicated.

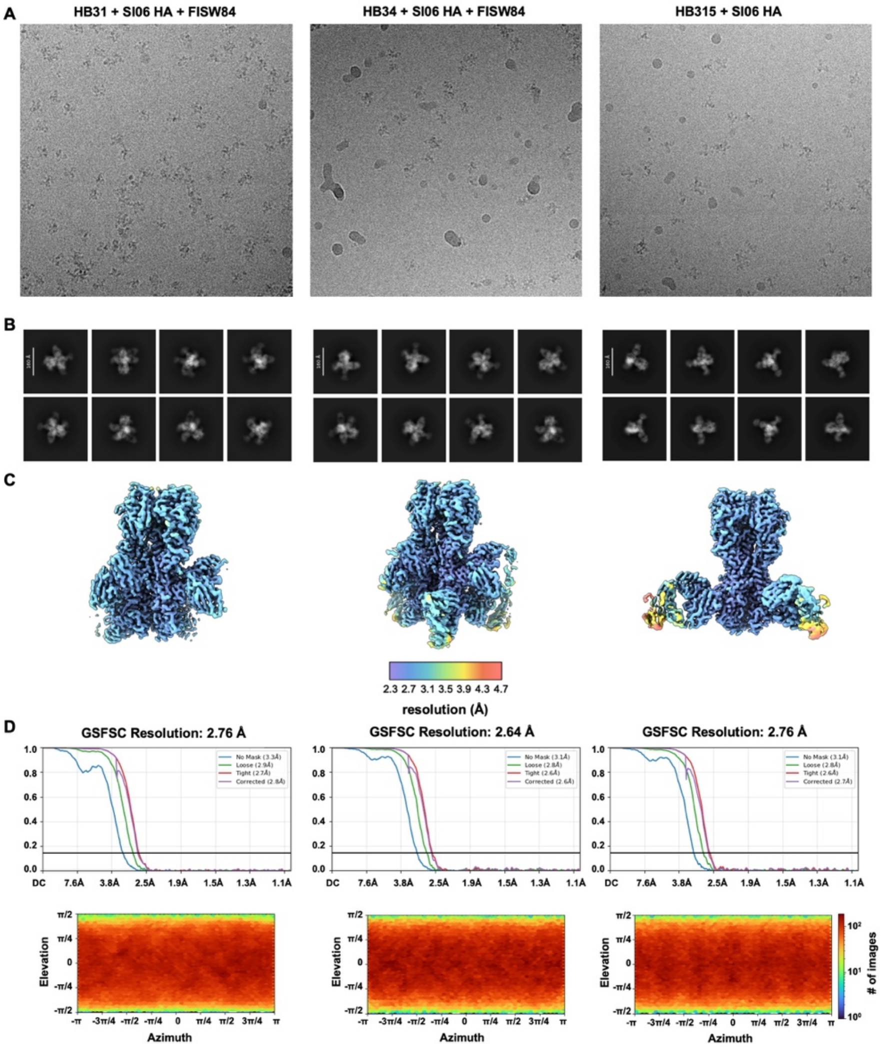


**Figure S7. Cryo-EM data processing for the structures of HB31, HB34 and HB315 Fabs in complex HA. (A)** Representative micrographs, **(B)** representative 2D class averages, **(C)** local resolution maps and **(D)** Fourier shell correlation (FSC) curves are shown.

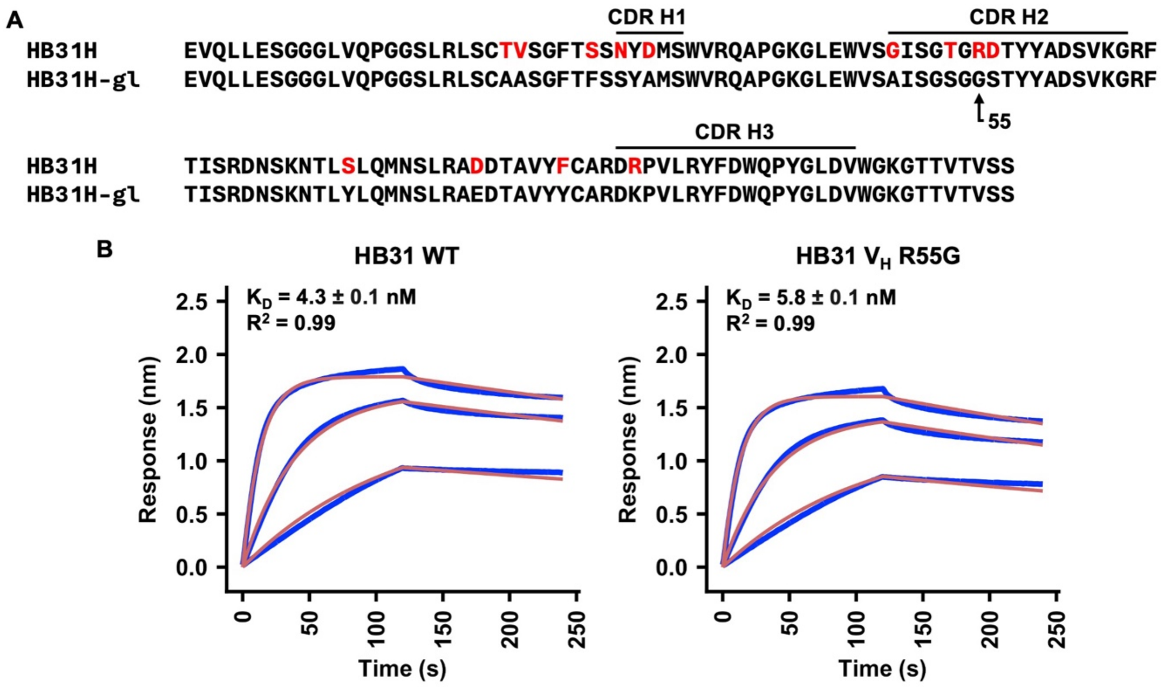


**Figure S8. Sequence determinants for the upper HA stem pocket of HB31. (A)** Sequence alignment of HB31 heavy-chain variable domain with its inferred germline sequence was performed using MAFFT[6]. Residues that represent somatic hypermutations in the V gene are colored in red. **(B)** Binding kinetics of HB31 wild-type (WT) and $V_H$ R55G germline revertant against H1 stem were measured by biolayer interferometry (BLI). Y-axis represents the response. Blue and red lines represent the response curve and the 1:1 binding model, respectively. Binding kinetics were measured for 300 nM, 100 nM, and 33 nM of Fab. The dissociation constants ($K_D$) and the goodness-of-fit values ($R^2$) are shown. Dissociation constants (KD) are shown as mean ± standard deviation.

### SUPPLEMENTAL REFERENCES


1 Dreyfus, C. *et al.* Highly conserved protective epitopes on influenza B viruses. *Science* **337**, 1343-1348 (2012). https://doi.org/10.1126/science.1222908

2 Benton, D. J. *et al.* Influenza hemagglutinin membrane anchor. *Proc Natl Acad Sci U S A* **115**, 10112-10117 (2018). https://doi.org/10.1073/pnas.1810927115

3 Corti, D. *et al.* A neutralizing antibody selected from plasma cells that binds to group 1 and group 2 influenza A hemagglutinins. *Science* **333**, 850-856 (2011). https://doi.org/10.1126/science.1205669

4 Kallewaard, N. L. *et al.* Structure and function analysis of an antibody recognizing all influenza A subtypes. *Cell* **166**, 596-608 (2016). https://doi.org/10.1016/j.cell.2016.05.073

5 Wu, N. C. *et al.* Recurring and adaptable binding motifs in broadly neutralizing antibodies to influenza virus are encoded on the D3-9 segment of the Ig gene. *Cell Host Microbe* **24**, 569-578 e564 (2018). https://doi.org/10.1016/j.chom.2018.09.010

6 Katoh, K. & Standley, D. M. MAFFT multiple sequence alignment software version 7: improvements in performance and usability. *Mol Biol Evol* **30**, 772-780 (2013). https://doi.org/10.1093/molbev/mst010
